# Antibiotic perturbation of the human gut phageome preserves its individuality and promotes blooms of virulent phages

**DOI:** 10.1101/2025.02.07.636470

**Authors:** Eugen Pfeifer, Camille d’Humières, Quentin Lamy-Besnier, Florian Plaza Oñate, Remi Denisé, Sara Dion, Bénédicte Condamine, Marie Touchon, Laurence Ma, Charles Burdet, France Mentre, Erick Denamur, Eduardo PC Rocha, the PrediRes study group

## Abstract

The use of antibiotics disrupts the gut microbiota, potentially leading to long-term health issues and the spread of resistance. To investigate the impact of antibiotics on phage populations, we followed 22 healthy individuals two weeks before and up to six months after a three-day course of 3^rd^-generation cephalosporins. The populations of phages very rarely encoded antibiotic resistance genes and were mostly temperate including many phage-plasmids. Gut phages remained individual-specific even after microbiome perturbation by antibiotics. Yet, we found a 20% decline in phage diversity the day after treatment, alongside blooms of a few (mostly virulent) phages. We suggest that these temporary dominant phages contribute to the recovery of gut bacterial diversity through kill-the-winner dynamics. This is supported by the finding that several phages targeted *Parabacteroides distasonis*, a bacterium thriving after cephalosporin treatment, and which only proliferated when these phages were absent.Our findings suggest that phages play a crucial role in the gut microbiota’s response to antibiotics by contributing to the restoration of microbial balance and diversity.

**Highlights:**

- In healthy individuals, cephalosporin antibiotics cause an approx. 20% decline in gut phage richness that is restored after 30 days
- Human gut phage communities are specific of each individual, which is retained post-antibiotic treatment
- Antibiotic perturbation causes an increase in the number of dominant virulent phages supporting kill-the-winner dynamics as mechanisms for shaping bacterial diversity
- Dominance of *Parabacteroides distasonis* (after cephalosporin treatment) is likely disrupted through its specific phages

**eTOC:** Antibiotic-induced disruption of the gut microbiome leads to a transient loss of microbiome diversity (bacterial and viral) and causes blooms of bacteria and their viruses. Our analysis suggests that these viruses prey on dominant bacteria, prevent their dominance, and contribute to the recovery of microbiome balance.

**Graphical abstract:** 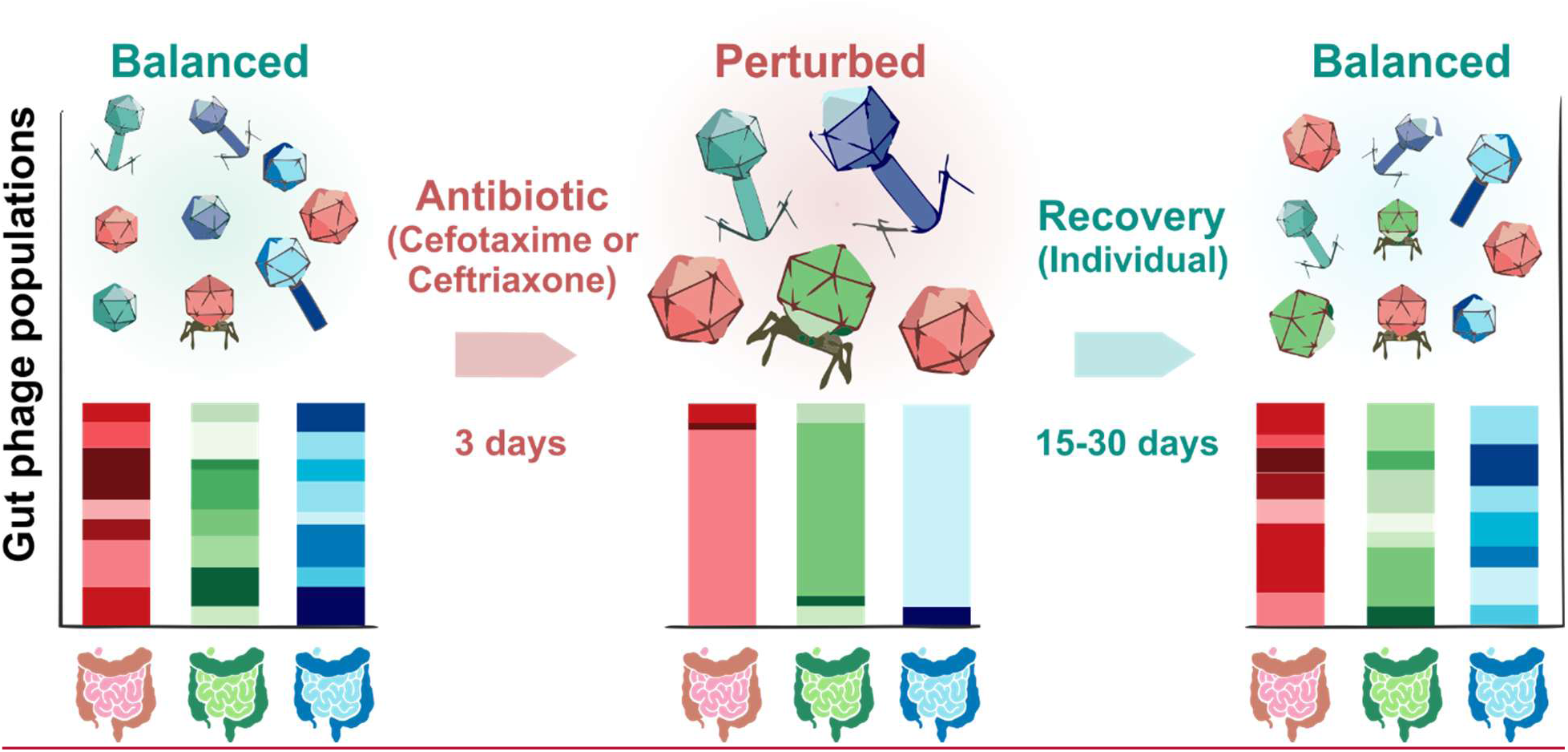

## Introduction

Changes in the gut microbiome’s composition frequently act as risk factors for the emergence of diseases, such as the proliferation of opportunistic microbes that thrive when the microbiome is destabilized^1^. The gut environment plays a crucial role in the emergence of antibiotic resistance genes (ARGs), as horizontal gene transfer within it facilitates their spread^2–4^. This process is further accelerated by the use of antibiotics favoring the growth of ARG-carrying bacteria. Gut commensals like *Candidatus Borkfalkia ceftriaxoniphila* or *Parabacterium distasonis* are known to bloom after treatment with cephalosporins^5^. Some species also release enzymes protecting vulnerable groups, as shown for gut commensal *Bacteroides spp*, who produce cephalosporinases that enhance the survival of susceptible community members from lethal doses of cefotaxime^6^. These dynamics lead to changes in the resistome (sum of ARGs within a microbiome) and can cause an unbalanced gut microbiota (microbial dysbiosis), which is commonly linked to gut-related diseases^7,8^. Antibiotics may be present in the gut when they are taken orally or when they are excreted through the digestive tract^9^. This is the case of ceftriaxone and cefotaxime, both 3^rd^ generation cephalosporins, which upon injection in patients are eliminated partly through the intestinal route and partly by the urinary tract^10,11^. The fraction of the antibiotics that is excreted by the digestive tract is subject to strong inter-individual variability, which is directly related with the impact of the antibiotic treatment on gut bacteria^12^. It is therefore essential to detail the impact of antibiotics on the human gut microbiome to better understand their effects on human health and on the spread of antibiotic resistance.

Bacterial viruses (phages) are highly abundant in the human gut microbiome and exhibit significant inter-individual variation within the intestinal tract^13^. It is suggested that they drive bacterial population diversity and dynamics ^14,15^. Some phages are virulent, they infect, replicate and lyse their hosts. Temperate phages can either behave similarly or become prophages, where they are kept in bacterial genomes (lysogen). Most prophage genes are silent, but some may provide adaptive traits to the lysogen. Some temperate phages propagate as phage-plasmids (P-Ps), i.e. they transfer vertically in the cell lineage as plasmids and horizontally as phages. We recently showed that P-Ps are widespread in bacteria^16^, and discussions on their ecological roles are gaining momentum^17^. Within the gut phageome (collective of phages), temperate phages were reported to be highly prevalent, with their numbers exceeding those of virulent phages^18,19^. In addition, it is estimated that 70-80% of gut bacteria carry prophages within their genomes^20,21^, suggesting that temperate phages have key roles in the gut environment. The impact of phages may be affected by antibiotics in multiple ways. First, antibiotics may induce temperate phages and result in lysis of bacterial populations. This has been reported for DNA damaging drugs such as fluoroquinolones^22,23^. Second, the change in bacterial populations following antibiotic therapy has almost necessarily an impact on their phages (which prey on them).

Finally, while ARGs are typically spread by conjugative plasmids requiring cell-to-cell contact, they are being increasingly identified in P-Ps that can transfer them and induce resistance by lysogenic conversion^24^, raising the possibility that P-P lysogens may protect bacteria from antibiotics. Yet, most past studies focused primarily on the impact of antibiotics on gut bacteria, and few aimed to understand the effects on the phages in the human gut. In particular, significant alterations of phage populations in the gut have been observed during antibiotic treatment for *Helicobacter pylori*, involving a proton pump inhibitor, amoxicillin, and clarithromycin^25^. Furthermore, a pioneering study including ten healthy volunteers, who received four antibiotics (quinolones, β-lactams, tetracyclines, and macrolides), suggested an expansion of ARGs in phages, although phages with ARGs remained at low frequency^26,27^. A general large-scale analysis of the impact of clinically-relevant antibiotics on the phage populations and on how this can feedback on the bacterial fraction is missing.

To improve our understanding of antibiotic-induced perturbation effects on the gut microbiome, we conducted the CEREMI clinical trial^28,29^, where ceftriaxone and cefotaxime were given intravenously over three days (clinical standards) to 22 healthy volunteers. Samples were taken 14 days before (day −14) and up to 180 days (day 180) following treatment, including before (day −1) and after (day 4) treatment (see Figure S1 for detailed sampling scheme). Stool samples were analyzed using measures of antibiotic concentrations, metabolomics and also metagenomics of bacteria, fungi and phages^28^. Samples before antibiotic treatment were used as controls since the strong inter-individual variability of the microbiome (bacteria and phages^13,30^) means that using other individuals provides a weak control. In all but two cases, the antibiotics were not detectable in the stool samples, suggesting β-lactam degradation by gut anaerobes, as previously described^31^. Furthermore, we observed a marked increase in the abundance of genes encoding β-lactamases in the bacterial fraction, and we detected disturbances in population structures (including richness) of both bacteria and phages following treatment^28^.

Our previous analysis focused on a broad comparison among the multiple variables of the study (bacteria, resistome, fungal composition, phage, measure of beta-lactamases, metabolomics, etc), leaving no space to detail the analysis of gut phages. Furthermore, the techniques used at the time hampered the study of the phage fraction (limited coverage of read signal). Hence, we have completely re-analysed the sequencing data to tackle questions that remained unanswered: i) How strong and how targeted is the treatment’s impact on the gut phageome? ii) Do gut phages carry β-lactam resistance genes, potentially contributing to their dissemination? iii) If so, are P-Ps involved and how common are they? iv) Finally, how does the treatment alter phage population dynamics? To take on these questions, we conducted a comprehensive, in-depth analysis of the phageome dataset^28^, applying state-of-the-art assembly techniques and metagenomic tools to expand and annotate the repertoire of gut phages. We searched for P-Ps by comparing the updated phageome data with P-P genomes^32^, employed a unified gut phageome database, and annotated ARGs using the NCBI AMRFinderPlus pipeline^33^. This enabled us to track changes in the populations of phages in terms of richness, diversity, relative abundances, and gene context throughout the surveillance period.

## Results

### Many novel high-quality genomes detected in the CEREMI dataset

Given the great advances in assembling and annotating phage metagenomics data in the last few years, we completely re-analysed the CEREMI phageome data set using two different assembly strategies (see Methods). We employed the most commonly used and recent programs (geNomad^34^, viralVerify^35^ and VIBRANT^36^) to fetch contigs classified as viral by at least one method (Figure S2AB). We then further recovered phage genomes with lower coverage by mapping all the reads on the large catalog of human gut virome datasets, the Unified Human Gut Virome (UHGV) database. We pooled all viral sequences to define dereplicated, viral operational taxonomic units (vOTUs). This allowed us to obtain a much more comprehensive dataset that resulted in a 2.6-fold increase (relative to our previous study^28^) in viral sequences to 6,467 unique vOTUs (Table S1), most of which originated from UHGV (Figure S2B). We found only 31 species of viruses infecting eukaryotes which were present at low relative frequency.

Hence, we will refer to vOTUs as phage species interchangeably throughout this study. The majority of sequences (84.8%) exceeded 10 kb in size (Figure 1A, Table S1). The sequence size distribution and the quality metrics of our dataset indicate a dataset with quality similar to that of UHGV (Figure S3AB), even if in our dataset all sequences ≥3 kb, regardless of the quality were included (UHGV used sequences >3 kb only if they had at least 50% completeness and, otherwise, a 10 kb cutoff was applied). Among the vOTUs, 796 were classified as complete by CheckV^37^ and 1150 as high-quality, accounting for 30.1% of the total (Figure 1B). The comparison of the vOTUs with all sequences from UHGV suggests that 1,951 are novel species (Figure S3CD). Although sequences of novel species were more likely to fail the CheckV^37^ quality control (15.6% versus 5.8% for the entire dataset, Table S1, Figure 1B, Figure S3C), this may in part stem from the difficulty of the tool in classifying novel sequences. A viral taxonomy was assigned for the majority of the vOTUs (98.1%), of which 98.5% are tailed dsDNA phages (*Caudoviricetes*). Only 73 vOTUs are ssDNA phages (ino- and microviruses) (Table S1).

**Figure 1:**
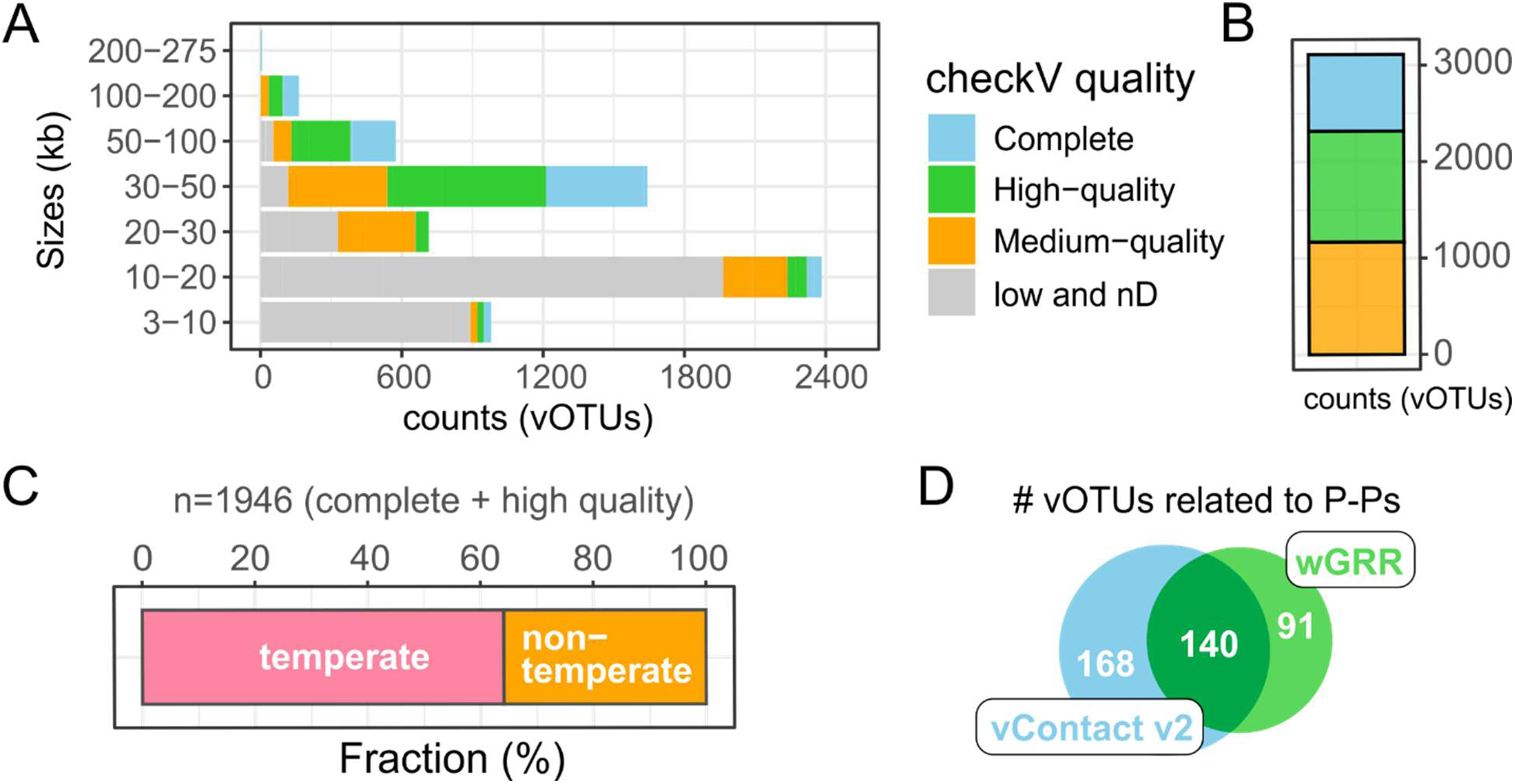
Overview of the gut phageome retrieved from 22 healthy donors. A. Sizes of all viral contigs (vOTUs), ranging from 3 kb to 275 kb, along with quality information (from CheckV^37^). B. Quality assessment of viral contigs was performed using CheckV^37^, with counts of medium, high-quality, and complete sequences shown (n=3,113). C. Lifecycles of 1,946 vOTUs (complete and high quality, assessed by CheckV^37^) were predicted using BACPHLIP^40^. D. Number of vOTUs related to P-Ps from a database of our previous work^32^. We used two methods, vConTACT v2^46^ and gene repertoire relatedness (wGRR), that employ shared protein networks to compute genome-to-genome relatedness (see Methods).

To exhaustively allocate the reads to the genomes, we sequentially mapped them against: the vOTUs, assemblies classified as non-viral, integrated prophages (identified by geNomad, VIBRANT, and CheckV), the UHGV, and the Unified Human Gastrointestinal Genome (UHGG) database (see Methods). Notably, we used the UHGG database to identify read sequences originating from bacteria, as it comprises over 200,000 non-redundant genomes from more than 4,644 gut prokaryotic species^38^. On average, 57.8% of the reads mapped to the vOTUs (Figure S4, Table S2), 18.9% aligned to sequences that could not be classified as viral, likely representing bacterial or highly divergent viral sequences, and only 0.32% mapped to prophages suggesting that their contribution to the read signal is minor. Around 4.58% of the reads mapped to viral contigs from UHGV that we did not include in our analysis due to very low sequence coverage. Finally, 11.8% of the reads mapped to sequences from UHGG, potentially representing bacterial contamination or highly diverse prophages not yet identified or represented in the databases used in this study. Interestingly, we identified six vOTUs with exceptionally large genome sizes (Table S1). These included five phages (>200 kb^39^) and a novel member of *Megaviricetes* (vOTU_5310), a class of eukaryotic giant viruses.

To predict bacterial hosts, we retained predictions for vOTUs retrieved from the UHGV, and for the others, we used iPHoP, resulting in assigning a host taxonomy to 5,378 vOTUs (Table S1). The hosts were found among the taxonomic classes of bacteria detected in the bacterial fractions^28^ with a preponderance of *Clostridia*, *Bacteroidia*, and *Bacilli* in similar ratios as those of UHGV (host of viral sequences) and of UHGG (bacterial sequences) (Figure S5A). However, the bacterial fractions revealed a higher prevalence of *Clostridia* and a lower prevalence of *Bacteroidia* species. We compared these values with other studies summarized in UHGG and found that the variance of the reported ratio *Bacteroidia*/*Clostridia* is high and that a few studies found even lower frequencies of *Bacteroidia* (Figure S5B).

### Temperate phages including phage-plasmids are prevalent in the gut

We applied BACPHLIP^40^ to predict the lifecycle of phages having complete and high-quality genomes (predicted by CheckV^37^, n=1946). The majority was classified as temperate (64.1%, Figure 1E). We then searched specifically for P-Ps, which are also temperate phages but typically lack integrases and are often misclassified by tools predicting phage lifecycles. We grouped our database of 1416 P-Ps^32^ with all vOTUs and used protein-sharing networks for the clustering (see Methods). We could thus identify 399 vOTUs that grouped with P-Ps representing 6.2% of all vOTUs (Figure 1F). Nearly all (87.2%) of these vOTUs clustered with putative P-Ps infecting Firmicutes and Actinobacteria, including *Bacillus thuringiensis, Clostridium beijerinckii*, *Corynebacterium atypicum*, *Clostridium perfringens* and *Enterococcus faecium* (Table S3, supplementary file 1). In particular, 198 clustered with a putative P-P from *B. thuringiensis* CTC, an episomal element described to be a 25-kb linear plasmid with similarities to a phage of *Lactococcus lactis*^41^. 61 vOTUs are similar to P-Ps of *C. beijerinckii* and/or *E. faecium,* that are described as linear elements with sizes of 16 and 18 kb, respectively. Among these, ϕ6423 (of *C. beijerinckii*) is suggested to be a phage^42^ whereas no information on a lytic cycle is available for p63-3 of *E. faecium*. Many vOTUs (n=44) grouped with a putative 48-kb P-P from *C. atypicum,* for which a phage lifecycle has been proposed^43^, but confirmation is lacking. 32 vOTUs clustered with vB_CpeS-CP51, a temperate phage infecting *Clostridium perfringens*^44^, for which no experimental evidence of a plasmid-life cycle is reported. With *Carjivirus communis*, the prototypic crAssphage, we grouped 11 vOTUs. *C. communis*, was recently reported to propagate as a P-P in its host^45^.

### β-lactam antibiotics induce a strong perturbation in the gut phageome with a distinct response for each individual

We assessed changes induced by the cephalosporin antibiotics along time by monitoring the total species richness (Figure 2A). On average, we detected 374 different vOTUs per sample, but the individual values were widely variable ranging from 38 to 771 (Figure S6A). While 3487 vOTUs (94 viral clusters (VCs)) were unique to a single donor, seven vOTUs (9 VCs) were detected in 21 volunteers (Figure S6B, Table S4). Due to the high variability and specificity of phage species within individuals^13^, their numbers strongly correlate with the number of samples analyzed. We therefore assessed changes in phage richness by using similar numbers of samples per time point (N=19 to 20). We excluded days with a low number of samples (two samples of day −7, and two samples of day 90), and pooled donor-specific samples of day 4 (N=19) with day 7 (N=1) and day 15 (N=6) with day 30 (N=14) (see sampling scheme in Figure S1). We observed a drop of 20.4% in total phage richness following the beginning of antibiotic treatment (days 4-7), which is comparable to the drop in bacterial species richness (19.3%, obtained from the bacterial fractions^28^, Figure 2A). Since we do not expect cephalosporins to directly affect viral particles, the simultaneous reduction in phage and bacterial richness likely reflects the impact of the decrease in bacterial hosts following the treatment on the phage population.

**Figure 2:**
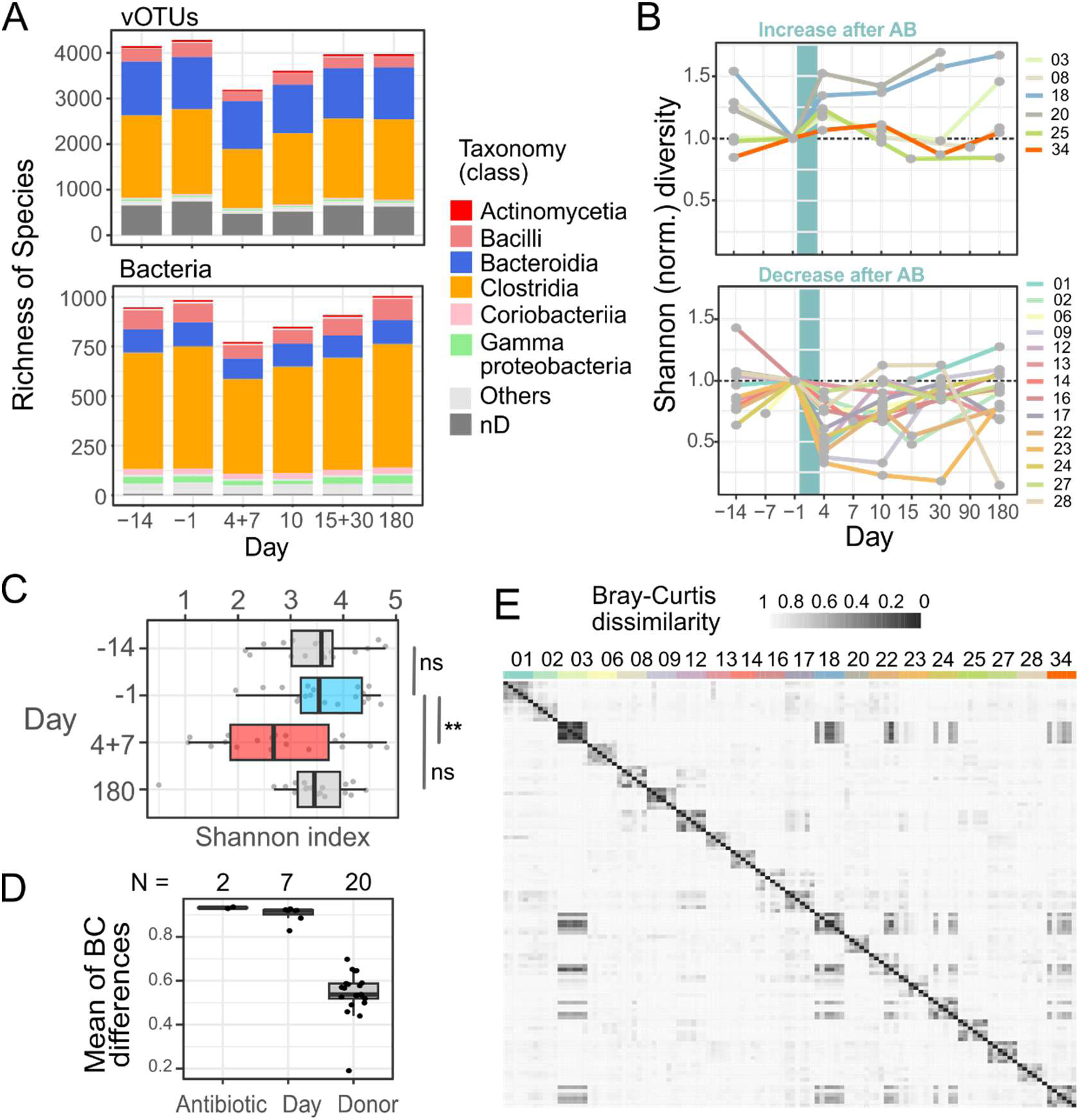
Perturbation effect assessed by total richness and samples’-diversities. A. Richness of vOTU and bacterial species per day of sampling. Not all volunteers provided samples on every collection day, which reduced species richness on certain days. To maintain a comparable sample size across time points, we pooled samples from day 4 with day 7, and from day 15 with day 30. For each of these pairs of days, volunteers provided a sample on only one of the two days. B. Shannon diversity index normalized to the day before treatment (day −1) for samples from all donors throughout the clinical trial (AB: antibiotics). The period of antibiotic treatment is indicated by a Cyan vertical bar. C. Shannon diversity of samples was assessed at days −14, −1 (blue, day before treatment), 4+7 (red, directly after treatment), and 180 post-treatment. Significant changes were determined using the Wilcoxon paired test, considering only donors with samples available for all these time points (N=18). **: p-value=0.007. D. Mean variances between samples based on Bray-Curtis distances, grouped by antibiotic type, sampling day, and donor. E. Matrix showing Bray-Curtis distances between samples (vOTUs), ordered by donor and within each donor ordered by sampling day.

To assess the strength and specificity of the perturbation, we compared compositional changes per volunteer over the trial. Only donors who gave samples on the day before treatment (day −1) were considered, excluding subject 11 and 29. We computed a Shannon index normalized per donor with respect to day −1. We observed major fluctuations before and after treatment, regardless of the type of antibiotics used (Figure S6C), with a higher number of donors whose diversity decreased in the days following treatment (N=14) when compared to those where it increased (N=6, Figure 2B). The phage populations experienced a substantial shift in diversity in some cases and a more moderate change in others. Globally, the treatment induced a significant decrease in diversity (Wilcoxon paired test, p = 0.006), which was more pronounced than the differences observed between pre-treatment samples (days −14 and −1) or between −14 and 180-days post-treatment samples (Figure 2C). After 180 days, in 15 out of 20 donors, similar levels of diversity were observed in comparison to compositions before treatment. These changes paralleled globally those of the bacterial population (Figure S6DE), where in 18 out of 20 donors, we observed a decline in bacterial diversity. Interestingly, in the two donors that showed increased bacterial diversity, we did not observe an increase in phage diversity, showing that bacterial and phage diversity are not always directly correlated.

To evaluate how the samples grouped by antibiotic type, time points, and donor, we computed the Bray-Curtis distance between all samples (Figure 2DE). This revealed lower distances between samples from the same donors (Mean_diff_ = 0.58) than when comparing samples across days (Mean_diff_ = 0.85) or antibiotics (Mean_diff_ = 0.93). This suggests that the overall individuality of each human individual^13^ outweighs the effects of time and antibiotic type.

### Cephalosporin treatment does not globally promote phages with resistance genes

In our previous work, we observed that the frequency of genes encoding β -lactamases increased after administration of the cephalosporins^28^. We suspected that spread of resistance genes might be facilitated by gut phages encoding ARGs. To detect them, we annotated all vOTUs with AMRfinderPlus^33^, and found nine phages with resistance genes (Figure S7). Among these phages, five were classified with high quality, four are temperate (none of which is classified as a P-P). Four encode resistance against β-lactams, three against tetracycline, one against vancomycin, and one against streptogramin antibiotics. All the four β-lactamases are classed as *cfxA* that encodes a class A β-lactamase that is often detected in *Bacteroides* species^47^. This matches the corresponding genera of the vOTU hosts that were predicted to be *Bacteroides* and *Prevotella*. Notably, we did not detect any genes encoding extended-spectrum β-lactamases typical for *Enterobacterales* in the phage sequences, even though these genes had been found in our previous work (in bacterial fractions)^28,29^.

We inspected the relative abundances of these vOTUs, focusing on those encoding β-lactam resistances and on their variation in frequency following administration of the antibiotics (Figure 3). The relative frequency remained quite constant with maximum levels being less than 0.6% in all samples. In 16 donors, no major changes were detected along the sampling period. vOTU_3934 was found in nearly all volunteers but usually at very low frequency (n=19, mean abundance 0.07% per donor and sample).

**Figure 3:**
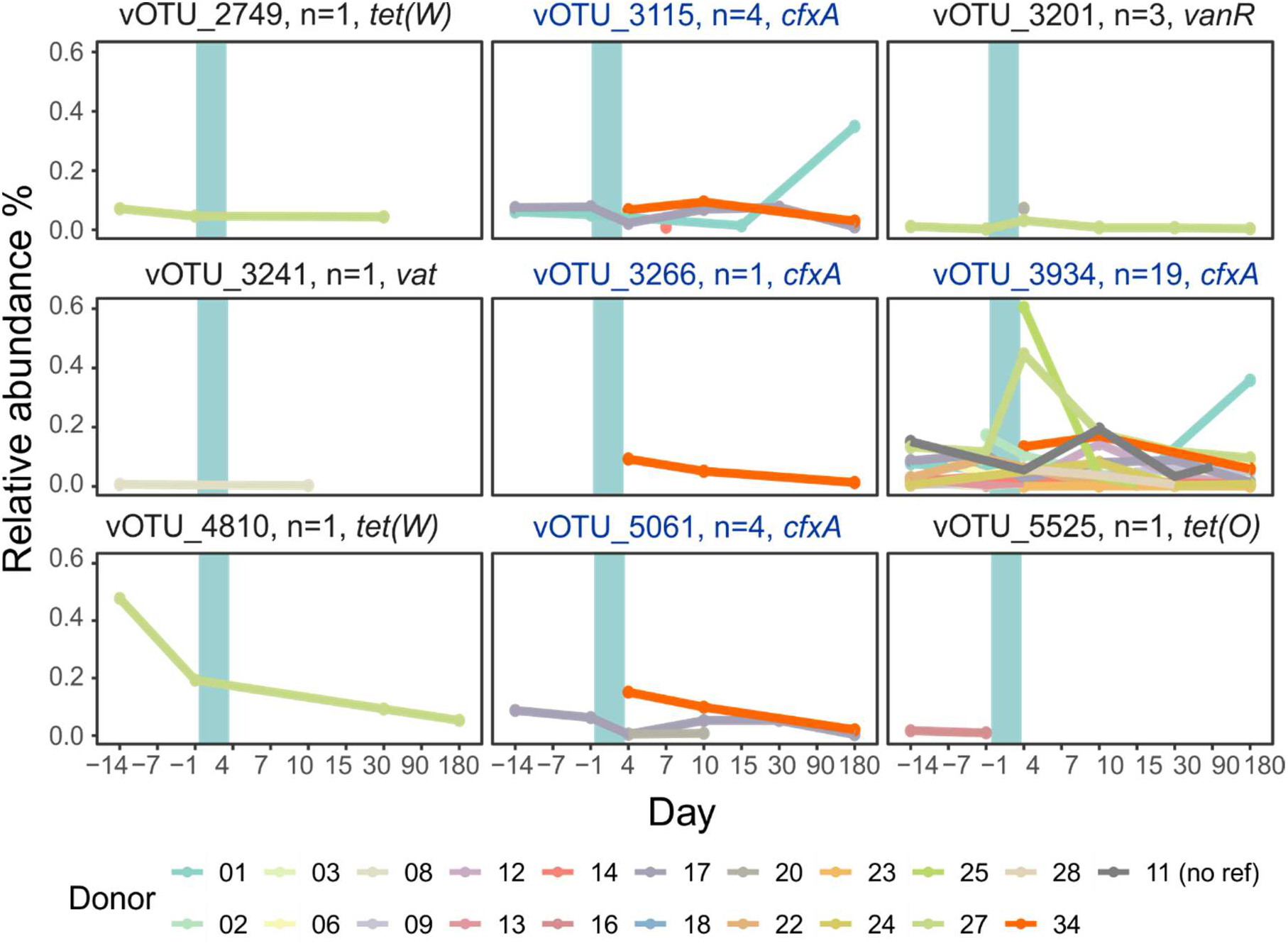
Relative abundance of vOTUs encoding ARGs. AMRfinderPlus detected nine vOTUs with genes encoding antibiotic resistance against: β-lactam (*cfxA*), tetracycline (*tet(O)* and *tet(W)*), vancomycin (*vanR*) and streptogramin (*vat*). N indicates the number of donors in which the vOTU was detected. The period of antibiotic treatment is indicated by the cyan bar. For donor 11, the reference sample from day −1 was not available.

However, in three donors (#34, #25 & #27) this phage increased in frequency after cephalosporin treatment. In donor #34 (orange in Figure 3), all four *cfxA* encoding vOTUs were detectable only on the day after treatment, and showed lower frequency after day 10 (post treatment). 180 days following the administration of antibiotics, we observed an increased frequency of vOTU_3115 and vOTU_3934 in donor #1 (shown in teal in Figure 3). This change is too late in the sampling period to be related to the use of antibiotics.

### Populations of phages undergo high fluctuations, with blooms after cephalosporin treatment

Our longitudinal study enables tracking the changes of phage species along time to assess the community dynamics. We quantified the relative abundances of all vOTUs and observed that phage populations fluctuate more widely than those of bacterial species as pointed out by higher coefficients of variation (CV) of relative abundances (Figure 4A, and Figure S8). On average, a vOTU had a relative abundance of 0.06%, and only a few species were highly abundant per sample (1% of all vOTUs with abundances >15% in at least one sample and 0.5% of vOTUs with abundances >25%) (Figure 4B, Table S5). We defined phages with quantities higher than 25% as dominant in the focal sample. We found 30 such phages (Figure 4A, Figure S9–S11). The dominant phages were detected in most donors, except in #08 and #34, albeit usually at much lower frequencies. Surprisingly, although most phages are predicted to be temperate, most dominant vOTUs were predicted to be virulent. In a few cases, the dominance was very high: four phages (vOTU_5377 infecting *Spirochaetia*, vOTU_6373 infecting *Prevotella*, vOTU_6374, and vOTU_6307) accounted each for more than 80% of the viral reads in the respective samples.

**Figure 4:**
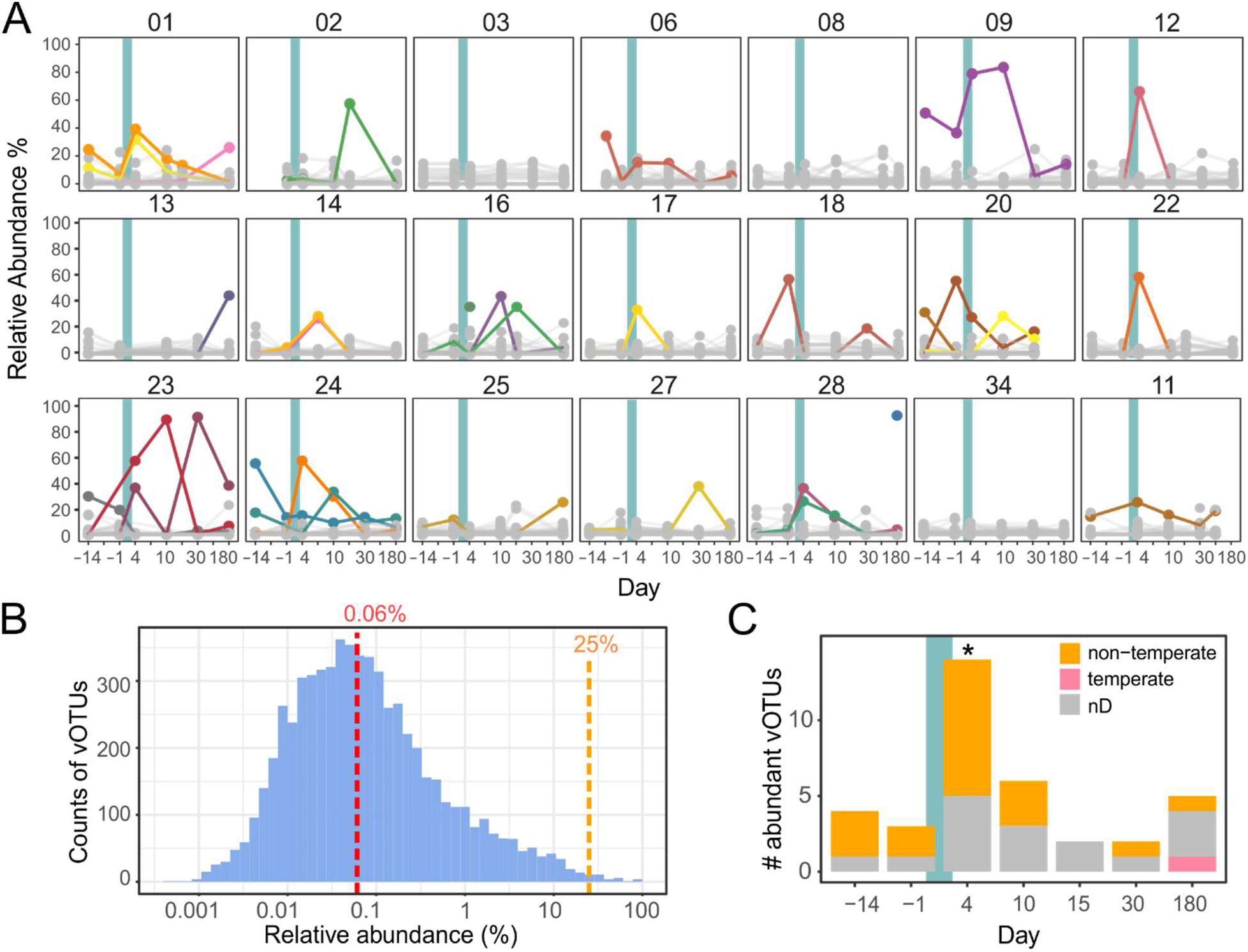
Dynamics of dominant gut phages. A. Shown are relative abundances of all vOTUs in all donor samples (N=21). The period of treatment with the cephalosporin antibiotics is indicated by cyan vertical bars. vOTUs were colored when their frequency exceeded 25%. For donor 11 no sample directly prior to treatment (day −001) was available. B. Histogram showing maximum abundance levels of all vOTUs. If a vOTU was detected in several donors, the highest level was considered. Average level is 0.06% (in red). The orange dashed vertical line indicate the threshold of 25% used to define the dominant species. C. Daily counts of dominant vOTUs (those with at least 25%) over all donors. As in A, the period of cephalosporin treatment is indicated by a cyan vertical bar. To determine significance, daily counts were compared to a Poisson distribution (see Methods), using the day −1 as the baseline. For day 4, a p-value of 0.012 was obtained (Figure S8 BC), indicating counts for this day are significantly different from the Poisson model.

Dominant phages undergo strong fluctuations in population sizes and only five dominant phages were found to persist across multiple time points in a few individuals. We tested if these fluctuations (counts of phages per day for the 30 dominant phages) follow a Poisson regression model and found that these events follow the expected distribution on most days, with the notable exception of the day after treatment (day 4). On this day, we detected a significant increase (p-value = 0.012, Figure S12AB) of the number of dominant phages (Figure 4C). This conclusion was robust to changing the threshold values for the dominant vOTUs in the regression analysis (Figure S12A). Overall, this indicates that the cephalosporin treatment affects the gut microbiota in a manner that results in spikes of high frequency of certain (mostly virulent) phages.

We did a similar analysis to the bacterial fraction obtained in our previous work, using the same threshold of 25% to detect dominant bacterial species. We identified 13 different bacterial species that became highly abundant in at least one sample (Figure S13, Table S6-S7), 10 of the class *Bacteroidia*, two of *Bacilli* and one of *Clostridia*. Of note, in seven individuals (#06, #11, #12, #13, #17, #18, #34), we did not detect any dominant bacterial species (according to our definition). Six out of the 13 species were found to be dominant across samples indicating that dominant gut bacteria tend to fluctuate less widely than highly abundant phages. In line with another work^5^, we observed that *P. distasonis* dominated the sample in donor #22, while in donor #16, we detected a high frequency (63.5%) of *Candidatus Borkfalkia sp003343765* post-treatment (Figure S13). Notably, another species of the same genus, *Candidatus B. ceftriaxoniphila*, has been reported to reach monodominance (abundances up to 92% after ceftriaxone administration)^5^. To explore the dynamics between these bacteria and their phages, we examined their co-occurrences. While we could detect phages of *Candidatus Borkfalkia sp003343765,* we found a few phages associated with *P. distasonis.* Interestingly, these phages became dominant in donors #01, #17, and #24, where *P. distasonis* has been detected but did not thrive (Figure 5). In contrast, in donor #22 (*P. distasonis* became dominant) none of the associated phages were detectable. These dynamics suggest that phages infecting *P. distasonis* may inhibit or substantially reduce its growth after cephalosporin treatment.

**Figure 5.**
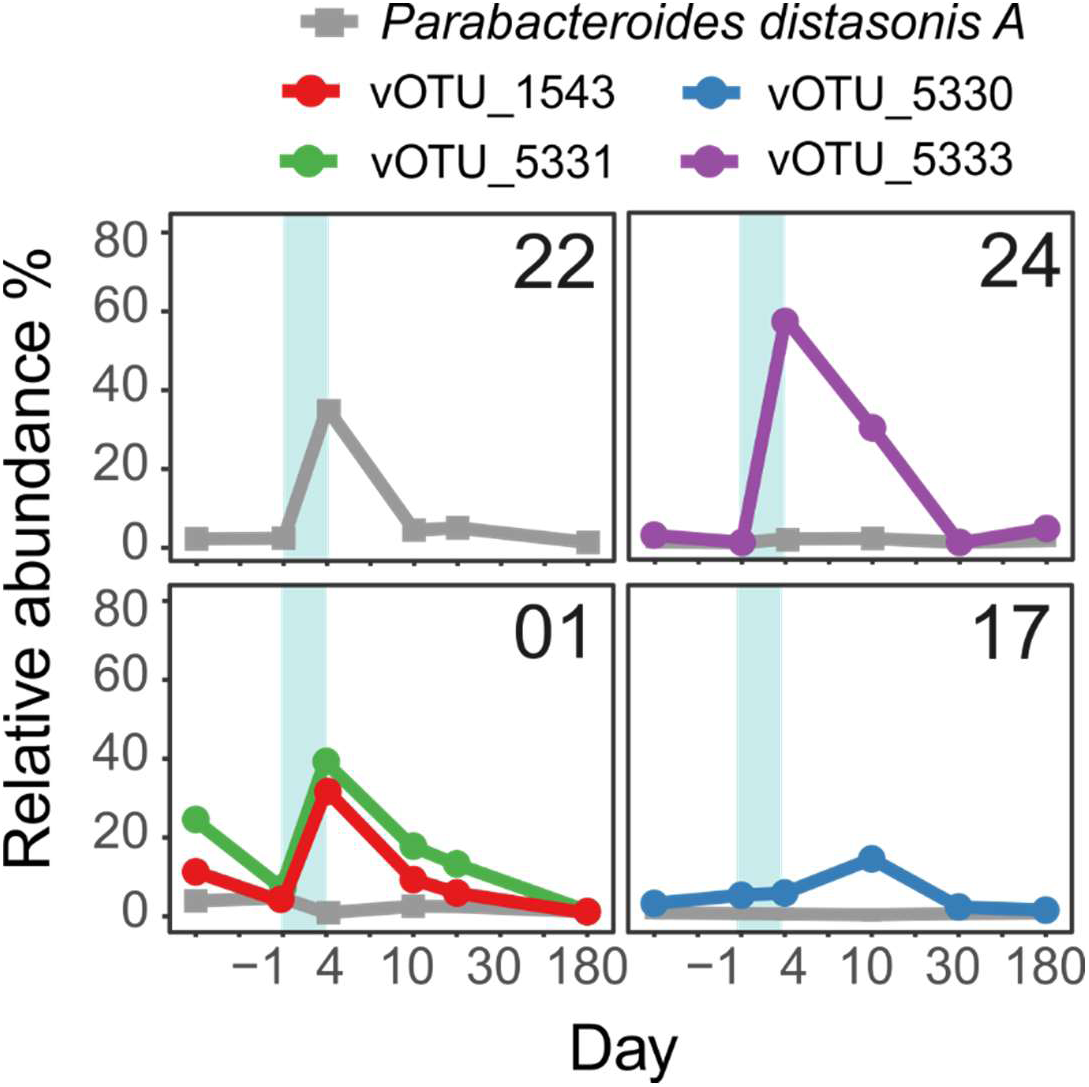
Abundance profile of *P. distasonis A* (grey) and its phages (purple, red, green, blue) in donor #01, #17, #22 and #24. Treatment by cephalosporin antibiotics are indicated by the cyan bar. All donors received ceftriaxone, except donor 17, who was treated with cefotaxime.

## Discussion

We aimed to understand the impact of antibiotics on the gut phage diversity by analyzing the phage fraction of healthy individuals receiving a standard antibiotic treatment with 3^rd^ generation cephalosporins. We used established protocols (fine-tuned in a past study^48^) and most recent computational resources, including the newest version of a harmonized human gut virome database, to recover a high quality dataset. Furthermore, we identified >1900 novel phage species, most of which were not closely related with any phage with a genome in GenBank. In consequence, 87.4% of the vOTUs could not be classified deeper than the class level (*Caudoviricetes*) and their bacterial host could only be assigned for roughly 79.5% (species level) of the vOTUs. These numbers are in line with those of other studies focusing on human gut phage diversity^19,49–51^, and fractions of prevalent hosts (*Bacteroidia*/*Clostridia*) aligned with distributions of large gut databases (UHGG and UHGV). However, we found a lower *Bacteroidia*/*Clostridia* ratio in the bacterial fractions in comparison to the viral hosts in viral fractions. The significance of this discrepancy remains uncertain, as studies with comparable bacterial species richness (n > 1160) report similar (lower and higher) distributions. This discrepancy was observed before and after treatment (Figure 2A) excluding a selective targeting of bacterial species by the antibiotics. Potentially, these differences arise due to intricate phage-bacteria dynamics. *Bacteroidia* species could be more susceptible to phage predation, resulting in their decreased levels. Furthermore, prophages are common in gut bacteria (70-80% lysogens, including *Bacteroidia* and *Clostridia*^20,21^), and polylysogeny (multiple prophages per genome) is highly prevalent ^52,53^. Consequently, the activation of various (pro)phage species within individual *Bacteroidia* genomes, yet poorly studied, could contribute to the observed differences.

We also found that phages exhibited larger fluctuations in frequency compared to bacterial species. We propose that several factors are affecting these dynamics. First, the higher diversity (richness) of phages compared to bacteria suggests intense competition for bacterial hosts, leading to shifting dominance where different phages succeed over time. Second, phages replicate quickly with high burst sizes, causing a rapid increase in population numbers under favorable conditions. Finally, on/off dynamics of prophages, especially in polylysogens, adds a further layer of high fluctuations.

We observed that the phage diversity is unique for the individual. This interindividual variation was the primary driver of variability between samples, consistent with findings in bacteria^30^ and another phageome study^13^. Here, we show that antibiotic-induced perturbation has a greater impact than longitudinal individual fluctuations, and that phage communities retain individual specificity post-treatment. We confirmed our previous results that the two cephalosporin antibiotics had similar impacts on the gut microbiome^28^, and revealed that the effect of the day of sampling, is much smaller than the effect of the donors. Hence, antibiotics may disturb the phage composition, but the latter remain largely a function of the individual.

In our previous studies, we observed an increase in genes encoding β-lactamases following cephalosporin treatment and isolated *E. coli* strains harboring β-lactamase genes^28,29^. We hypothesized that gut phages might contribute to the dissemination of these resistance genes. However, our analysis reveals that only a few gut phages encode β-lactamase genes. These phages appear to play a minor role in the microbial perturbation induced by antibiotics since they are found at low abundances even just after the end of the treatment when antibiotic impact is highest. Also, several of these phages are not ranked as high quality and could be other types of mobile genetic elements. Conservatively, only one out of the four identified vOTUs (vOTU_3115) ranks high in the characteristics of a phage providing resistance to bacteria: it has a high-quality genome of more than 30 kb (typical of tailed phages) with a high number of phage genes and is predicted to be temperate thereby potentially providing resistance by lysogenic conversion. Overall, this suggests that temperate phages carrying antibiotic resistance genes are rare and remain rare upon antibiotic treatment, presumably having minor effects in these individuals.

We found that temperate phages outnumber by almost 2 to 1 the virulent ones. This is in line with past work, detecting 76% of temperate phages in a Belgian cohort^18^ and 78% in A Danish cohort^19^, consistent with recent findings from a quantitative study on their numbers^53^. Interestingly, we observed a significant number of vOTUs (>6%) related to P-Ps, which appear to have been overlooked in previous gut phageome studies. The identification of P-Ps poses specific challenges since these are temperate phages that usually lack an integrase and there has been a paucity of methods to identify them in metagenomes. We opted for conservative methods, including gene sharing networks and stringent homology searches, to detect P-Ps. We avoided to predict P-Ps *de-novo* (as elements with genes typical of phages and plasmids), to minimize false positives that arise from the ambiguous classification of plasmids (as viral or chromosomal sequences^54^) or as results of recombination events. Consequently, our numbers of P-Ps in the gut environment are likely an underestimate. Many vOTUs clustered with P-P types that are experimentally poorly characterized. *C. communis* is one example, whose lysogenic lifecycle remains under debate. Experimental analysis of a recent study suggests that *C. communis* propagates primarily as a P-P^45^. Plaque formation has not yet been demonstrated (suggesting a lysogenic lifecycle), and genes typically associated with functions of integrases or an integrated state of the genome has not been observed^45^. Our search for plasmid replication genes identified vB_CpeS-CP51 (from *C. perfringens*) as a potential P-P. These genes are similar to those of *C. communis* (supporting a P-P lifecycle) and to ϕSM101 (P-P of *C. perfringens*^55^). While plasmid replication genes strongly suggest a P-P lifecycle, this classification remains tentative and requires experimental evidence. Further details on other P-Ps (putative and confirmed cases) that clustered with vOTUs of this study are described in Table S3. Our analysis highlights the prevalence of P-Ps among gut phages, likely driven by the high abundance of temperate phages. These results suggest that P-Ps play a significant role in ecological dynamics of gut phage communities, even as questions remain about their exact lifecycle mechanisms.

Although temperate phages are usually the most numerous, the phage species that we identified as becoming dominant following cephalosporin treatment are almost all predicted to be virulent. Some studies have reported an increase in virulent gut phages following perturbations of the gut microbiota. In a 1983 study, fecal samples were analyzed for phage content, revealing that lower titers from healthy individuals were predominantly composed of temperate phages, while higher titers from patients with intestinal diseases were mainly composed of virulent phages^56^. Similar findings were reported in a 2004 study, where stool samples from 140 children with severe gastroenteritis were analyzed for coliphages, and roughly 90% were identified as virulent T-even phages^57^. Nevertheless, the increase in virulent phages observed in our study was somewhat unexpected, given that β-lactam antibiotics are known to induce the bacterial SOS response in species like *E. coli*^58^, which can trigger the activation of prophages. However, the dominant phages are not predicted to be hosted by enterobacteria and we propose that the observed rise in virulent phages was driven through other circumstances. Specifically, we hypothesize that some of these phages prey on bacterial populations that temporarily became dominant after being stimulated by the antibiotic-induced perturbation. This hypothesis aligns with the ‘kill-the-winner’ model described for marine systems^59^, and which is hypothesized for the gut environment^60^. Since fast-growing bacteria are suppressed, this model predicts favorable growth of slow-growers, promoting their diversity^61^. Here, we present data for *P. distasonis* and its phages supporting this framework in multiple donors. We observed that high phage levels were correlated with low abundance of *P. distasonis*, while low phage levels coincided with a high abundance of the host (Figure 5). Species within the *Bacteroides* and *Parabacteroides* genera have increasingly developed resistance to certain antibiotics^62^, particularly *P. distasonis* isolates against β-lactams^63^. This increasing resistance contributes to their ability to thrive. We suggest that their phages could help to restore the diversity in the gut environment. By infecting and killing bacteria that are rising in frequency, they control the number of their hosts and regulate the bacterial populations that would otherwise have become, and potentially stay, dominant. By this, they create niches for other species to colonize the gut, thereby contributing to return to homeostasis after antibiotic therapy.

## Limitations of the study

Our study design is limited in that it provides only a snapshot analysis, allowing us to analyze phage frequencies only in relative terms and preventing us from capturing full dynamics over time. This could have implications in certain results, notably in the study of the dominant virulent phages as it could be argued that it is not these phages that become abundant, but all the others that became extremely rare. We find this hypothesis less parsimonious, because we showed that bacterial clades for which many phages were identified, e.g. the *Bacteroidia* and *Clostridia* (Figure 2A), were only moderately affected by the antibiotics, providing little reasons of why the frequency of their phages would systematically collapse. Additionally, given the high phage diversity, it is more reasonable to conclude that one phage became highly frequent rather than to suggest that nearly all other phages disappeared from the samples. Future studies need to focus on these specific interactions. In particular, in vitro experiments conducted in controlled environments, such as bioreactors, could provide valuable insights. By enabling precise cultivation and sampling under antibiotic exposure^64^, these experiments may significantly enhance our understanding of these dynamic interactions.

While our study suggests a limited and minor role for gut phages in transferring antibiotic resistance genes, this conclusion is based on data from healthy volunteers. Reports indicate that antibiotics can have lasting effects on the gut microbiota, including community shifts that favor the establishment of pathogens (*Clostridioides difficile*) and resistant bacteria^65,66^. It is plausible that patients with a history of antibiotic-resistant bacteria possess distinct phage populations with distinct genetic content and a higher propensity for transducing resistance genes.

Our results suggest that over 6% of the gut phage vOTUs were P-Ps. Detecting P-Ps is particularly challenging in metagenomic datasets because they are often split in multiple contigs. We clustered the detected vOTUs with a previously reported P-P database^32^. This approach fails to detect novel P-Ps because it is limited to the diversity of the known ones, and is also impeded by any misclassifications within that known set. While P-Ps are temperate phages, they do not integrate into the bacterial chromosome and typically lack integrases. Many tools designed to type viral life cycles, including BACPHLIP^40^, are trained on complete genomes and rely on typical phage genes involved in the lysogenic cycle as hallmarks (e.g. integrases and repressors). However, metagenome assemblies are often incomplete and may lack key functional modules, particularly those related to integration and lysogeny, leading to the misclassification of some phages, including P-Ps, as non-temperate.

## Resource availability

### Materials availability

CEREMI trial (ClinicalTrials.gov identifier <u>NCT02659033</u>)

### Data and code availability

The metagenomic shotgun sequencing data of bacterial microbiome are available from the European Nucleotide Archive under accession number PRJEB58157 (www.ebi.ac.uk/ena/browser/view/PRJEB58157). The metagenomic shotgun sequencing data of phage microbiome are available from the European Nucleotide Archive under accession number PRJEB58815 (www.ebi.ac.uk/ena/browser/view/PRJEB58815).

Rscripts, data tables, sequence and meta-information to reproduce figures and analysis are available on GitHub and Zenodo repositories under following links: https://github.com/EpfeiferNutri/PrediRes.git and 10.5281/zenodo.14758482.

### Consortia

Members of the PrediRes study group not otherwise listed as authors are listed here:

Xavier Duval (INSERM, Univ Paris Diderot, APHP-Bichat Hospital), Dusko Ehrlich (INRAE Metagenopolis), Laurie Alla (INRAE Metagenopolis), Emmanuelle Le Chatelier (INRAE Metagenopolis), Florence Levenez (INRAE Metagenopolis), Nathalie Galleron (INRAE Metagenopolis), Nicolas Pons (INRAE Metagenopolis), Benoît Quinquis (INRAE Metagenopolis), Khadija Bourabha (INSERM), Antoine Bridier Nahmias (INSERM, Univ Paris Diderot), Olivier Clermont (INSERM, Univ Paris Diderot), Mélanie Magnan (INSERM, Univ Paris Diderot), Dominique Rainteau (INSERM, Univ Pierre et Marie Curie, APHP–Saint Antoine Hospital), Antonin Lamazière (INSERM, Univ Pierre et Marie Curie, APHP–Saint Antoine Hospital), Emilie Gauliard (INSERM, Univ Pierre et Marie Curie, APHP–Saint Antoine Hospital), Farid Ichou (ICAN), Philippe Lesnik (ICAN), Marie Lhomme (ICAN). Jimmy Mullaert (INSERM, Univ Paris Diderot, APHP—Bichat Hospital), Thu Thuy Nguyen (INSERM).

## Acknowledgements

We gratefully acknowledge Marie-Agnes PETIT, and her team at MICALIS, for their valuable discussions and support.

This work was supported in part by the Laboratoire d’Excellence IBEID Integrative Biology of Emerging Infectious Diseases (grant ANR-10-LABX-62-IBEID), the PREDIRES project (grant ANR-16-CE15-0022), and the French junior professor chair ANR grant (ANR-22-CPJ1-0041-01).

High-throughput sequencing was performed on the Institut Pasteur Genomics Platform, member of “France Génomique” consortium (ANR10-INBS-09-08). This work used the computational and storage services (TARS cluster) provided by the IT department at Institut Pasteur, Paris.

## Author contributions

Conceptualization: EP, CdH, EPCR, ED

Biological analysis of the samples: CdH, SD

Computational analyses: EP, CdH, QLB, BC, FPO

Contributed material, software or expertise: MT, CB, RD, FPO

Sequencing of the samples: LM

Writing—original draft and designed figures: EP

Writing—review & editing: EP, CdH, ED, EPCR;

All authors revised and contributed to the current draft of the manuscript.

Supervision and funding acquisition: CB, FM, ED, EPCR.

## Declaration of generative AI and AI-assisted technologies

During the preparation of this work the authors used DeepL (DeepL SE, (2017)) and ChatGPT (OpenAI, (2021)) in order to improve the language and readability. After using this service, the authors reviewed and edited the content as needed and take full responsibility for the content of the publication.

## Supplemental information titles and legends

Supplementary file 1: Homology between phage-plasmids and vOTUs.

File contains comparison of methods that detected vOTUs as P-Ps (slide 2), summaries of vOTU and phage-plasmid networks based on wGRR (slide 3) and vConTACT v2 (slide 4). Synteny plots (generated with gggenomes, slide 5-32) show detailed comparisons of vOTUs and P-P genomes.

Table S1: Summary of vOTU characteristics.

Table S2: Read signal and its allocation on different gut microbiome related datasets.

Table S3: Summary of phage-plasmids to vOTU comparison.

Table S4: vOTUs across donors.

Table S5: Abundance profiles of vOTUs.

Table S6: Summary of microbial species.

Table S7: Relative abundance profile of microbial species.

## Supplemental figures

**Figure S1.**
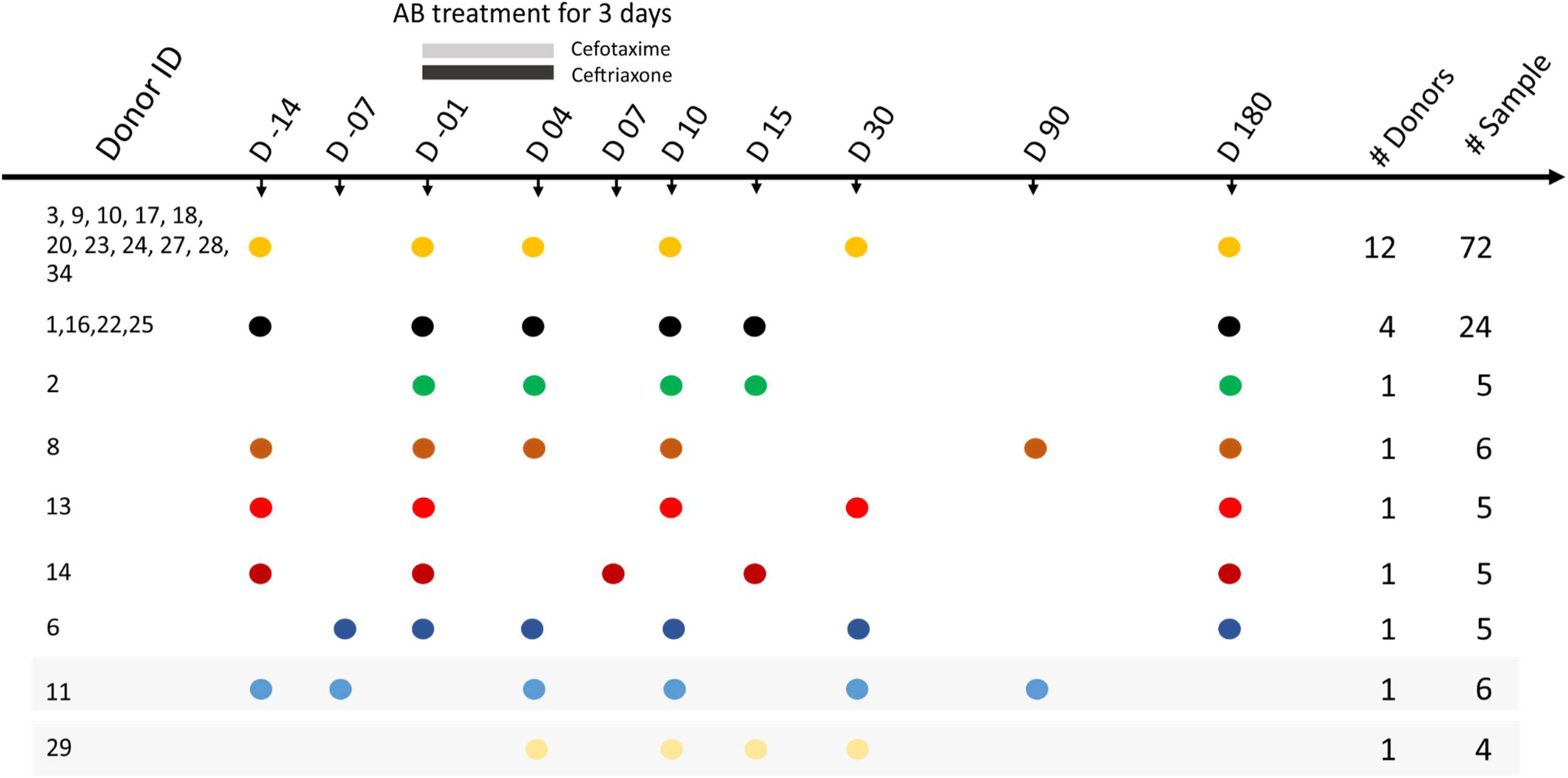
Detailed overview of sampling scheme and timeline of the clinical trial^28,29^. Samples from two donors (marked with grey bars) were partly excluded. Briefly, samples of donor 29 were excluded from ecological analyses (incl. diversity and abundance analysis) due to absence of any samples prior to the treatment. However, reads were included in the assembly and processing of viral contigs (see Methods). Donor 11 was excluded, if samples from the day −1 (before treatment) were required (specified in the analysis).

**Figure S2.**
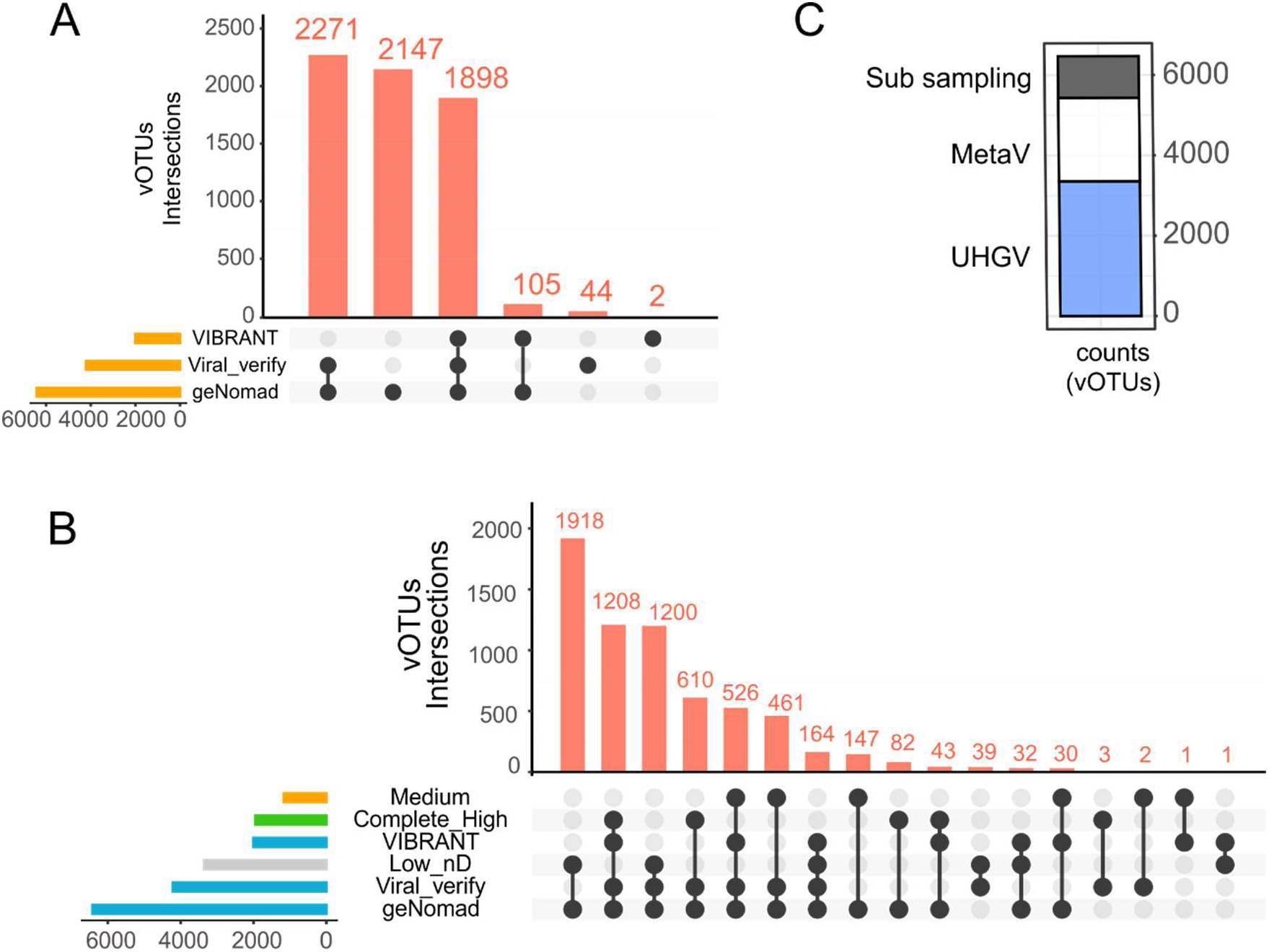
Recovery and quality of viral contigs (vOTUs). A: Counts of viral contigs detected by different classification tools (VIBRANT, viralVerify, geNomad). B: Quality of viral contigs (vOTUs), predicted by different tools, was validated by CheckV. Complete and High-quality, and Low-quality and not determinable vOTUs were count together. C. In total 6,467 vOTUs were recovered by mapping reads to the UHGV and following two assembly strategies (MetaV: MetaSPAdes --viral, Sub sampling: Fraction of reads with SPAdes, see Methods).

**Figure S3.**
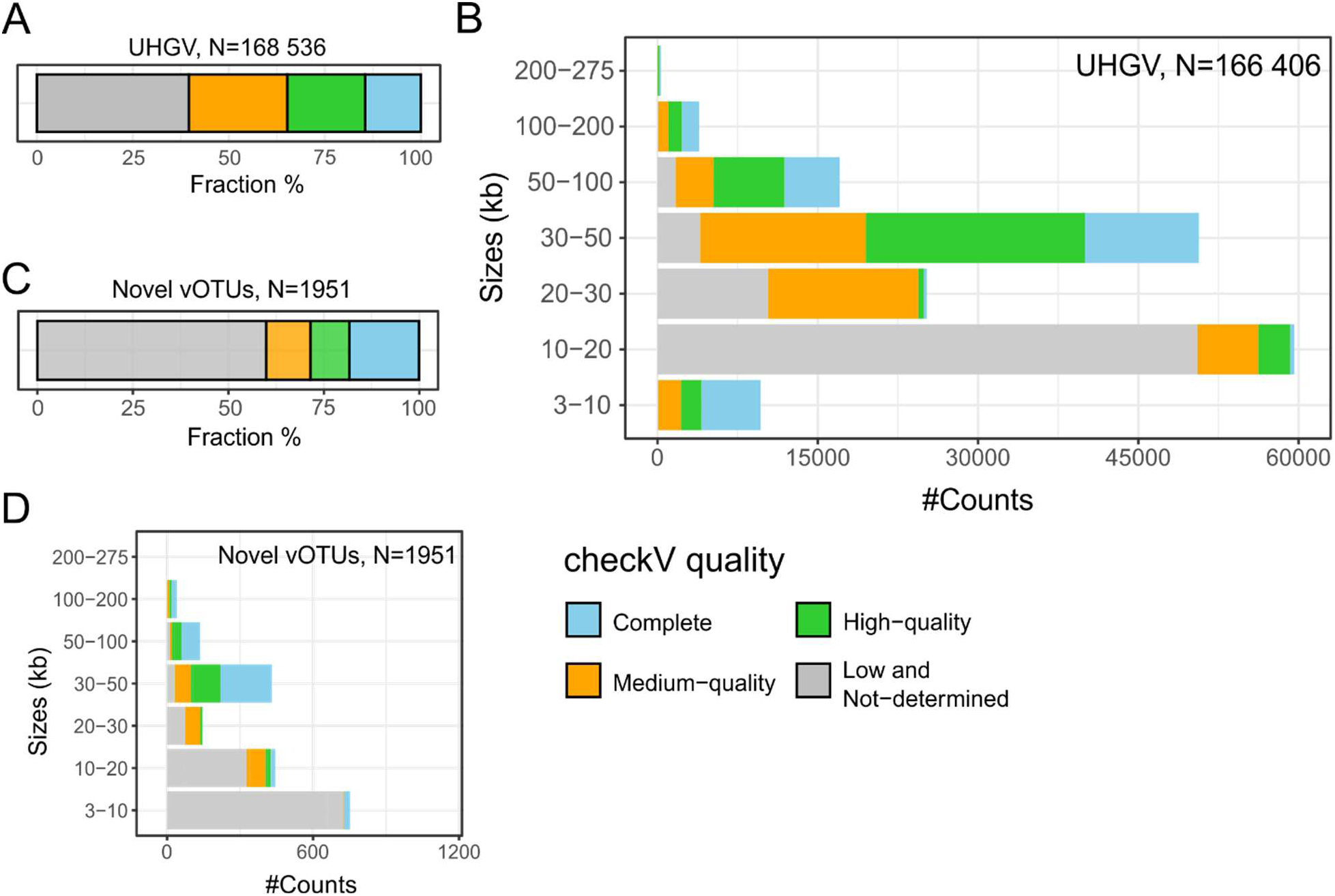
Completeness, quality, and sizes of sequences from UHGV and newly identified vOTUs. A and C: Quality assessment of sequences from UHGV (A) and of novel vOTUs (n=1,951) (C). B and D: Size distribution in relation to the quality of sequences from UHGV (B) and new vOTUs (D). Notably, UHGV only sequences >10 kb or with at least >50% completeness (medium quality) were considered (specified in https://github.com/snayfach/UHGV).

**Figure S4:**
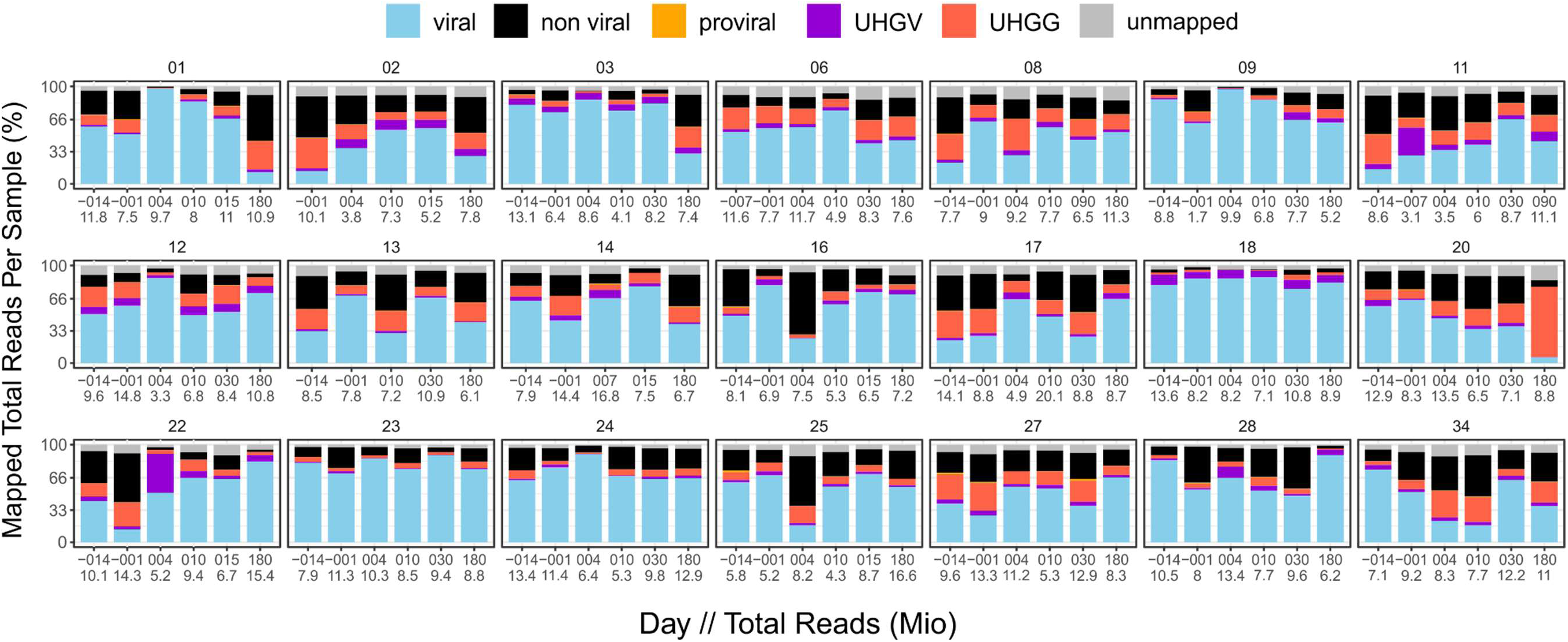
Allocation of the read signal to different gut-microbiome-related datasets. Reads from all samples were step-wise mapped on all vOTUs, non-viral assemblies, proviruses (predicted by geNomad^34^, CheckV^37^ and VIBRANT^36^), on the UHGV catalog and lastly on sequences of UHGG (see Methods). Fractions of the mapped signal are listed in Table S2. A. represents the average of all samples that are shown in bar plots B. In A, numbers below the boxes represent non-redundant sequences or representative genomes from the database. Numbers above the boxes represent the number of reads and fractions mapping on the datasets. In B, numbers below the bar plots indicate the sampling day and the number of reads in millions.

**Figure S5.**
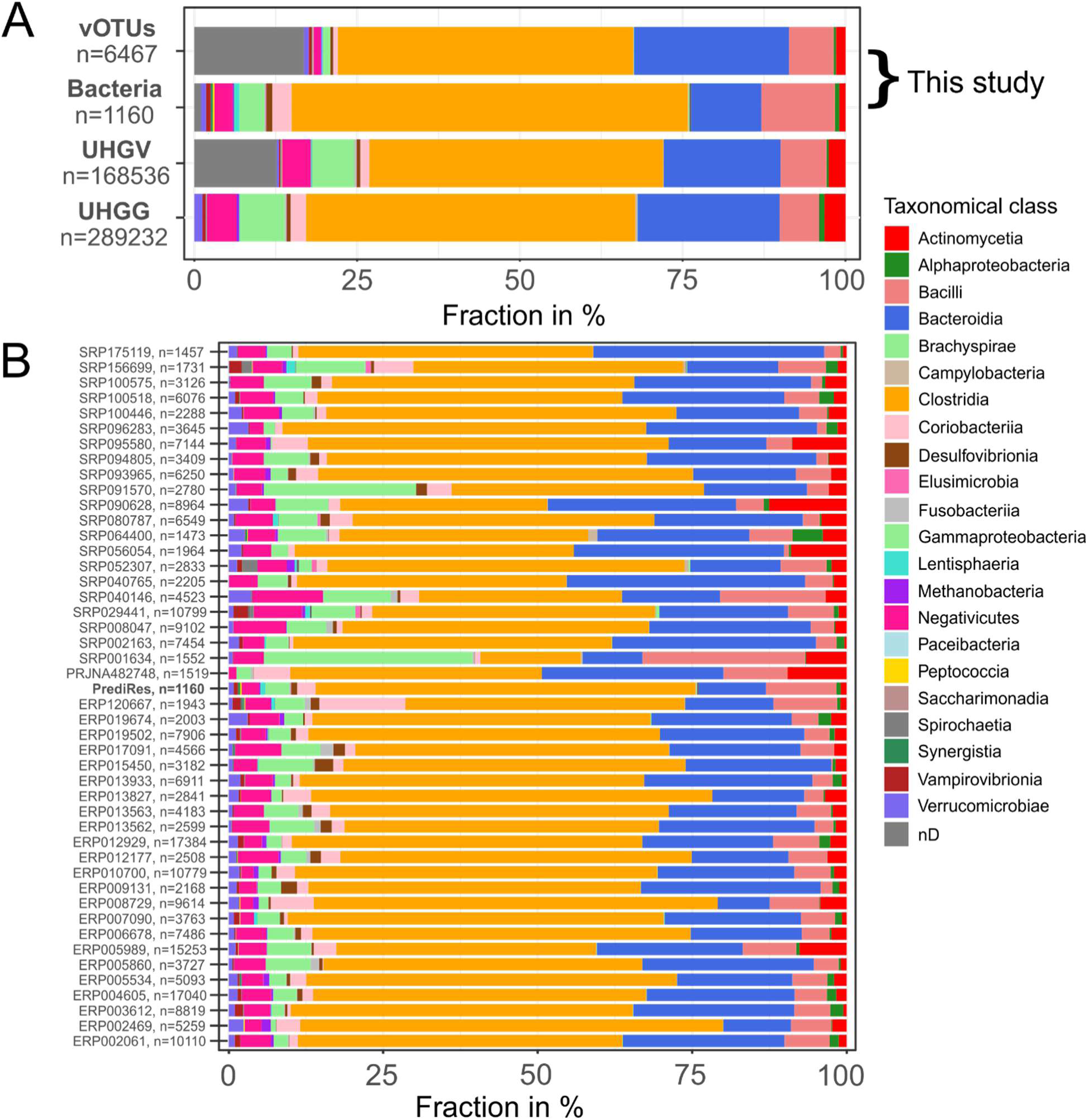
Taxa (class level) of hosts of the gut phages and of detected bacteria. A. Hosts of gut phage samples (vOTUs of this study and of UHGV), and bacterial species (of bacterial fractions of this study and of UHGG). B. Taxa of bacterial species detected in 47 studies including this work (PrediRes) with more than 1160 species.

**Figure S6.**
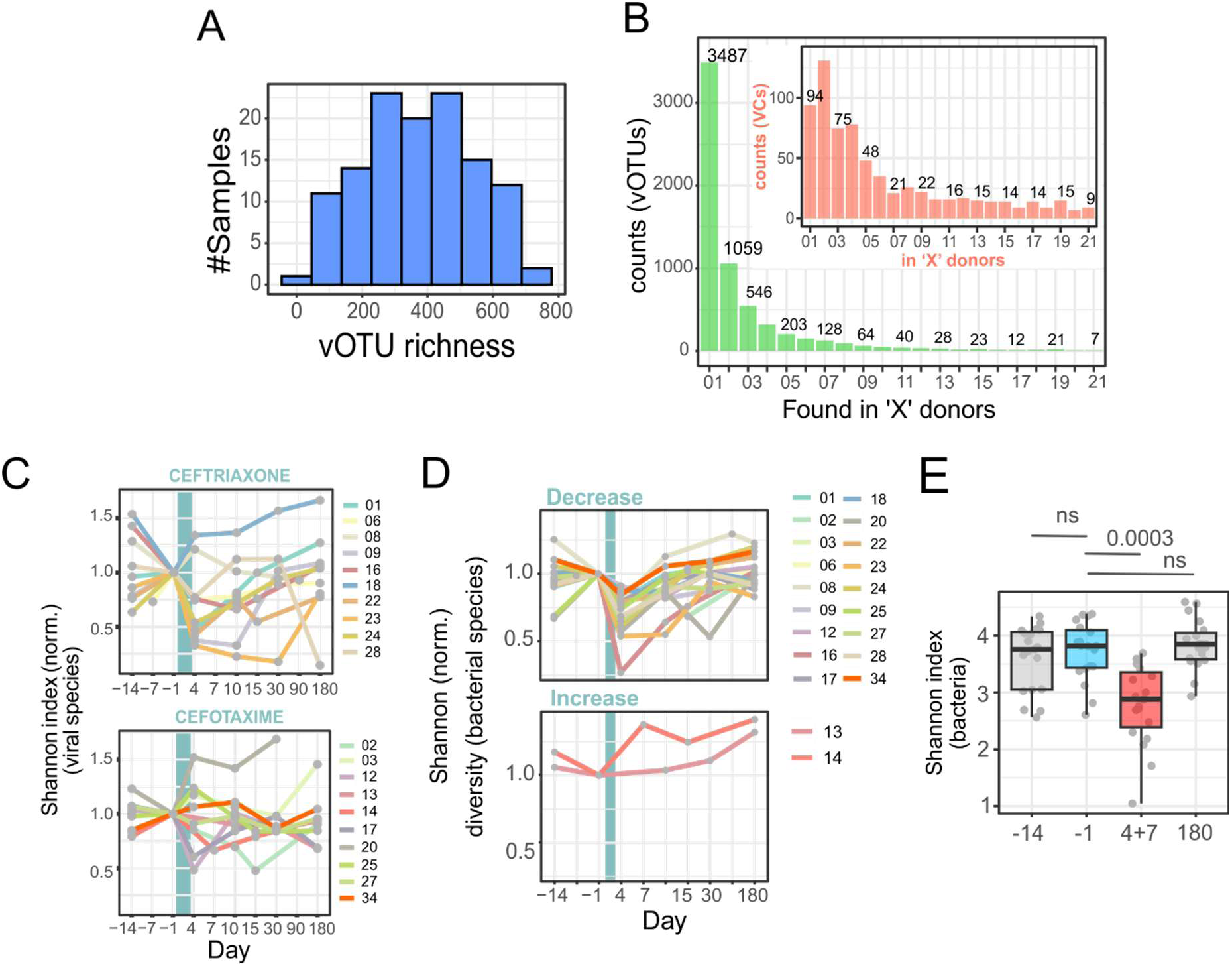
vOTUs and bacterial richness and diversity. A. Histogram displaying the range of richness per sample across a total of 123 samples. B. Counts of vOTUs (green bars) and viral clusters (VCs, red bars) across donors. C. Shannon diversity of vOTUs for the samples separated by type of cephalosporin. The period of antibiotic treatment is indicated by a Cyan vertical bar. D. Normalized diversity of the bacterial species. Abundances of bacterial species were computed from full metagenome samples (mostly bacterial species) and used to calculate the Shannon diversity per sample. This index was then normalized (per donor) to the day before treatment. The individuals were split in terms of decreasing or increasing diversity after treatment. E. As Figure 2C, but for bacterial species

**Figure S7.**
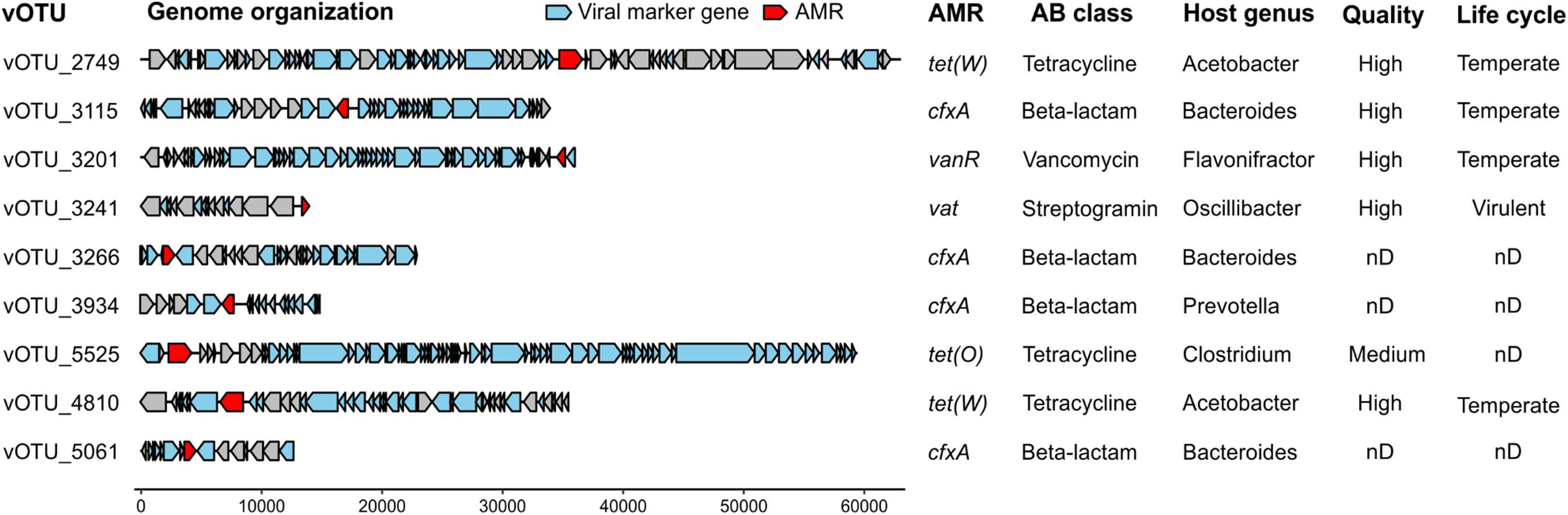
Genome organization of vOTUs encoding antibiotic resistance genes. Viral genes annotated by geNomad are shown in blue and ARGs by AMRFinderPlus in red. nD: not determinable.

**Figure S8.**
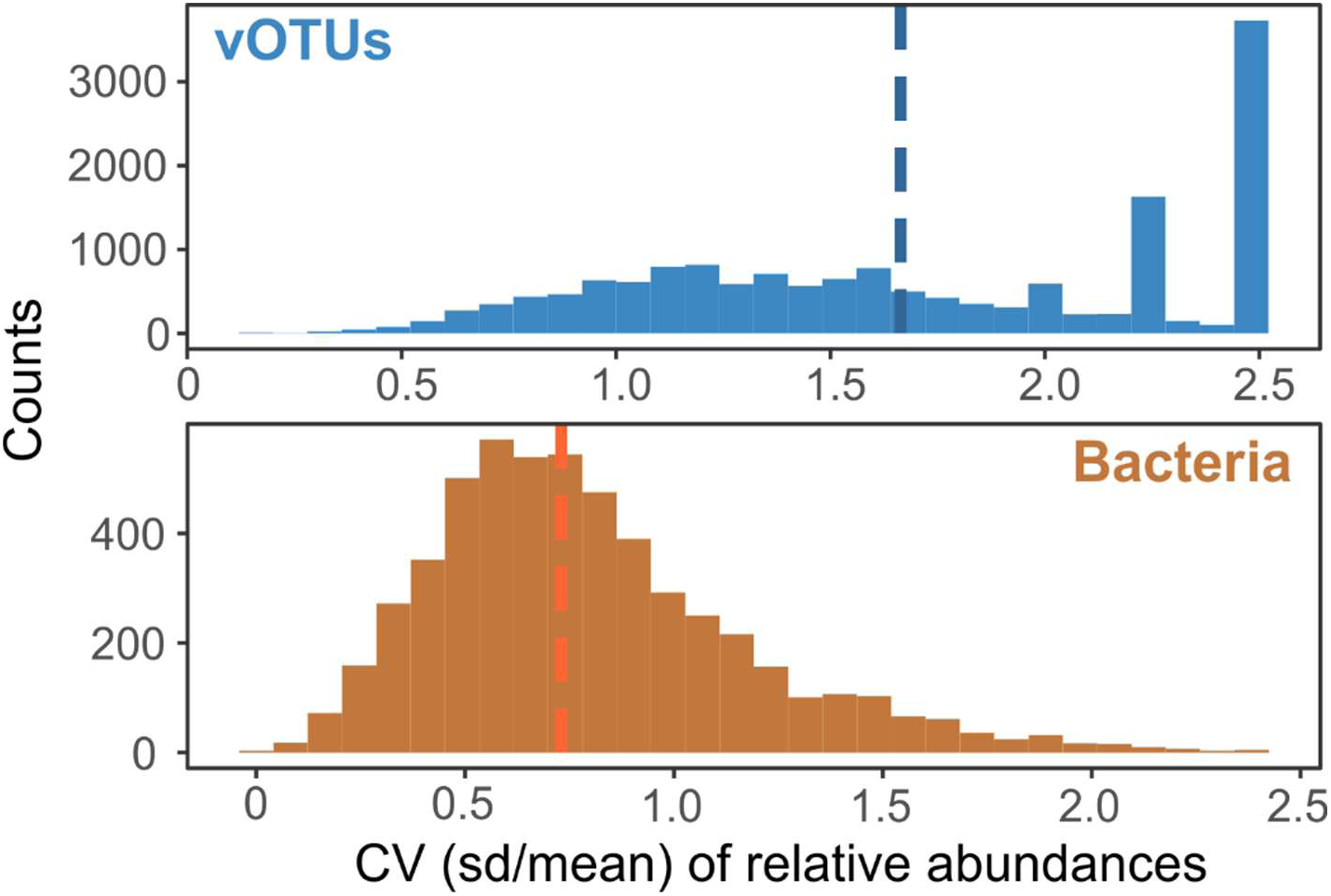
Relative standard deviation of abundance levels for phage and bacterial species. The coefficient of variation (CV) was calculated for all phage species (upper panel, blue) and bacterial species (lower panel, orange), with each CV based on a minimum of three values. Median values are indicated by dashed lines. Species absent in all samples from a donor were excluded from the analysis. For species detected at least once in a donor, absences were assigned a value of 0.

**Figure S9.**
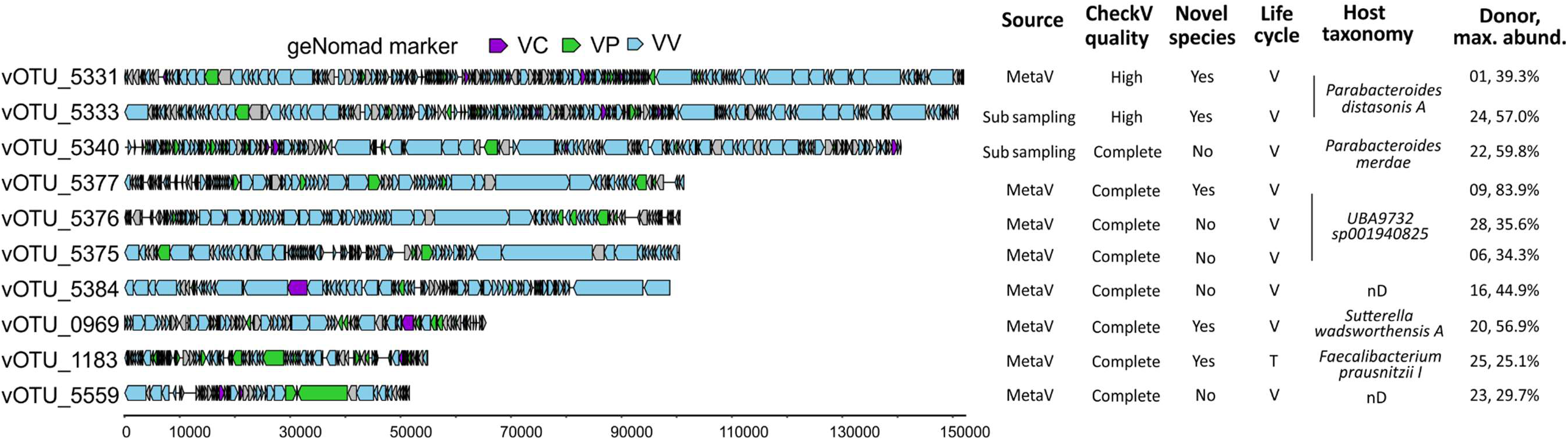
Genome organization of first 10 out of the 30 dominant vOTUs (max. abundance >25%, sorted by size). geNomad marker genes (two letter code: v=viral, p=plasmid, c=chromosomal), are colored. geNomad uses frequency of marker genes to classify sequences into viral, plasmids and chromosomal contigs (see https://portal.nersc.gov/genomad/marker_features.html). Life cycle V: virulent, T: temperate. nD: not determinable.

**Figure S10.**
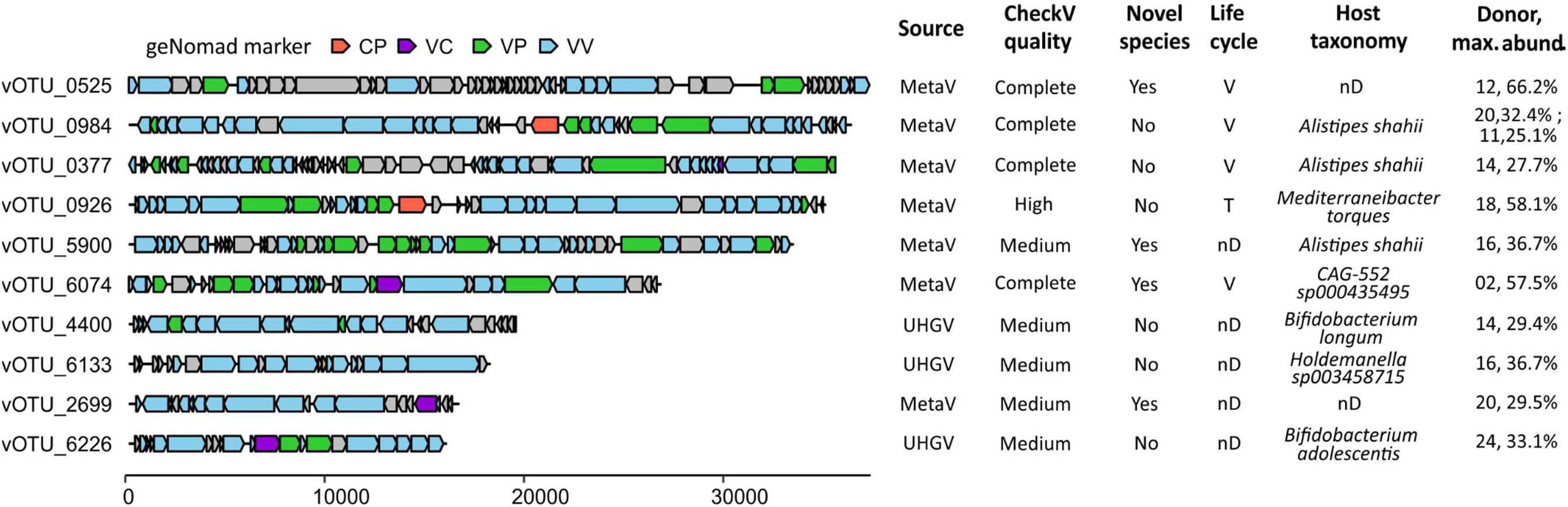
As Figure S9, but shown are 11 to 20 out of the 30 dominant vOTUs.

**Figure S11.**
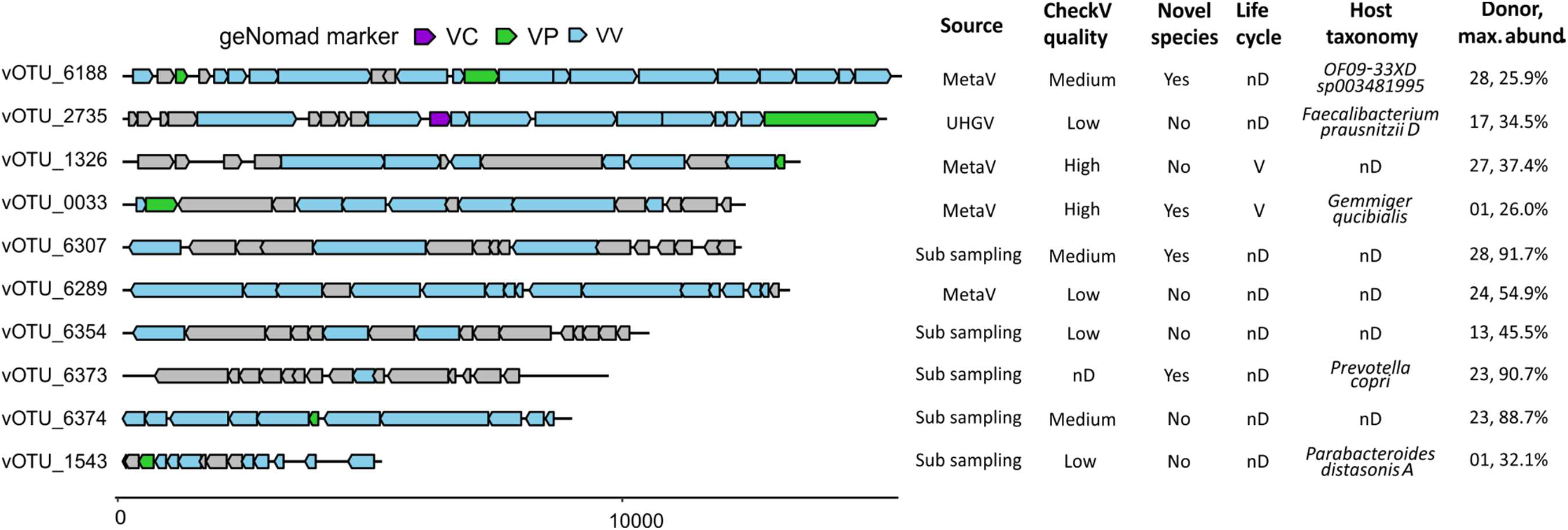
As Figures S9, but shown are 21 to 30 out of the 30 dominant vOTUs.

**Figure S12.**
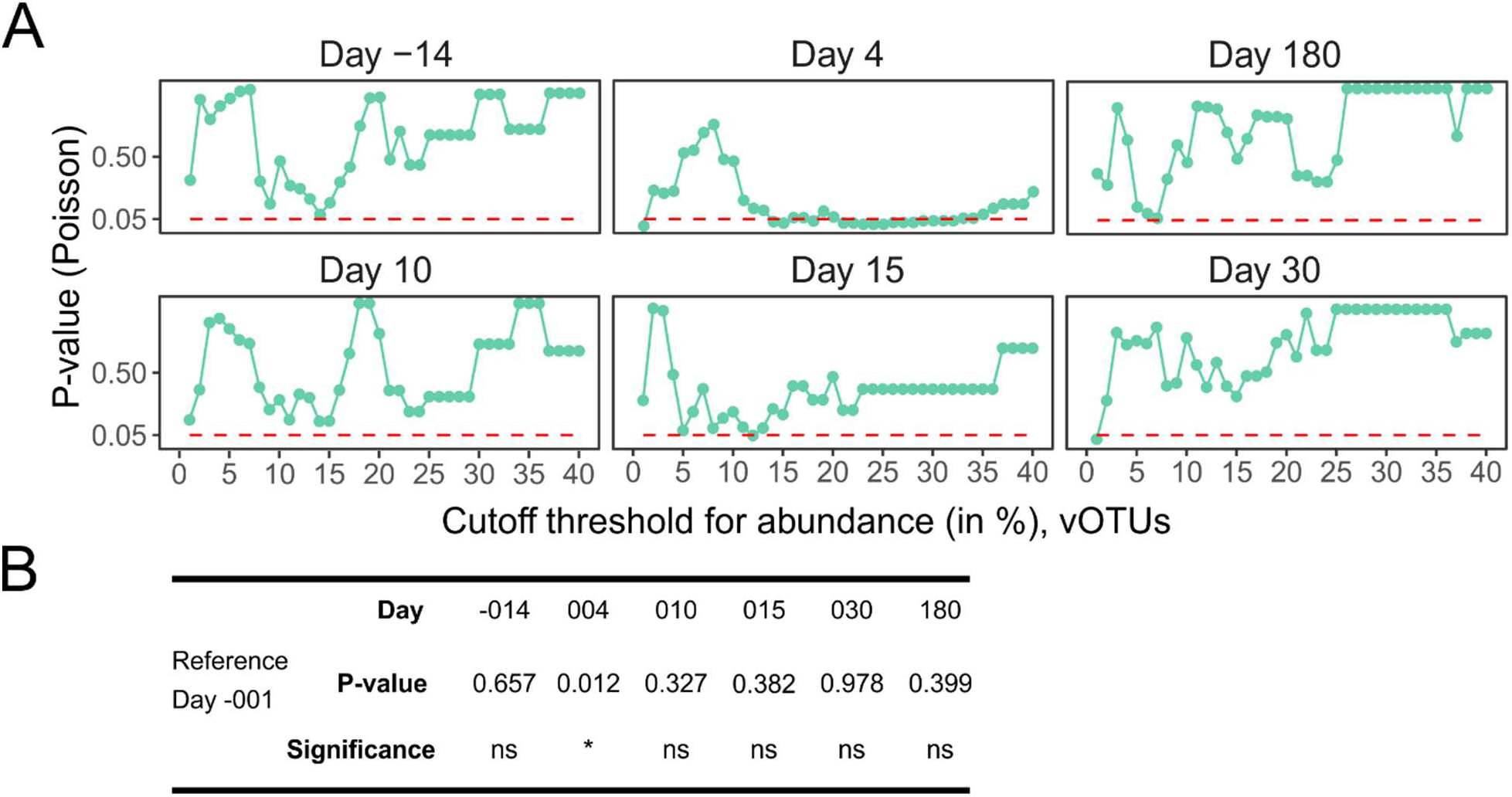
Poisson regression tests. A. The number of dominant phage species depends on the definition of dominance: in our study we used the threshold of 25%. We also tested various other values in the range 0.1% to 40% for defining dominance. At each cutoff, we counted the number of dominant species and tested if these events follow a Poisson distribution. The day before treatment (day -001), was set as the reference value in the GLM (see Methods). A p-value of <0.05 (dashed, horizontal red line) indicates these events are significantly different (do not follow the Poisson distribution). B. P-values computed at a cutoff of 25% abundance (to define dominant species) are shown (exemplary).

**Figure S13:**
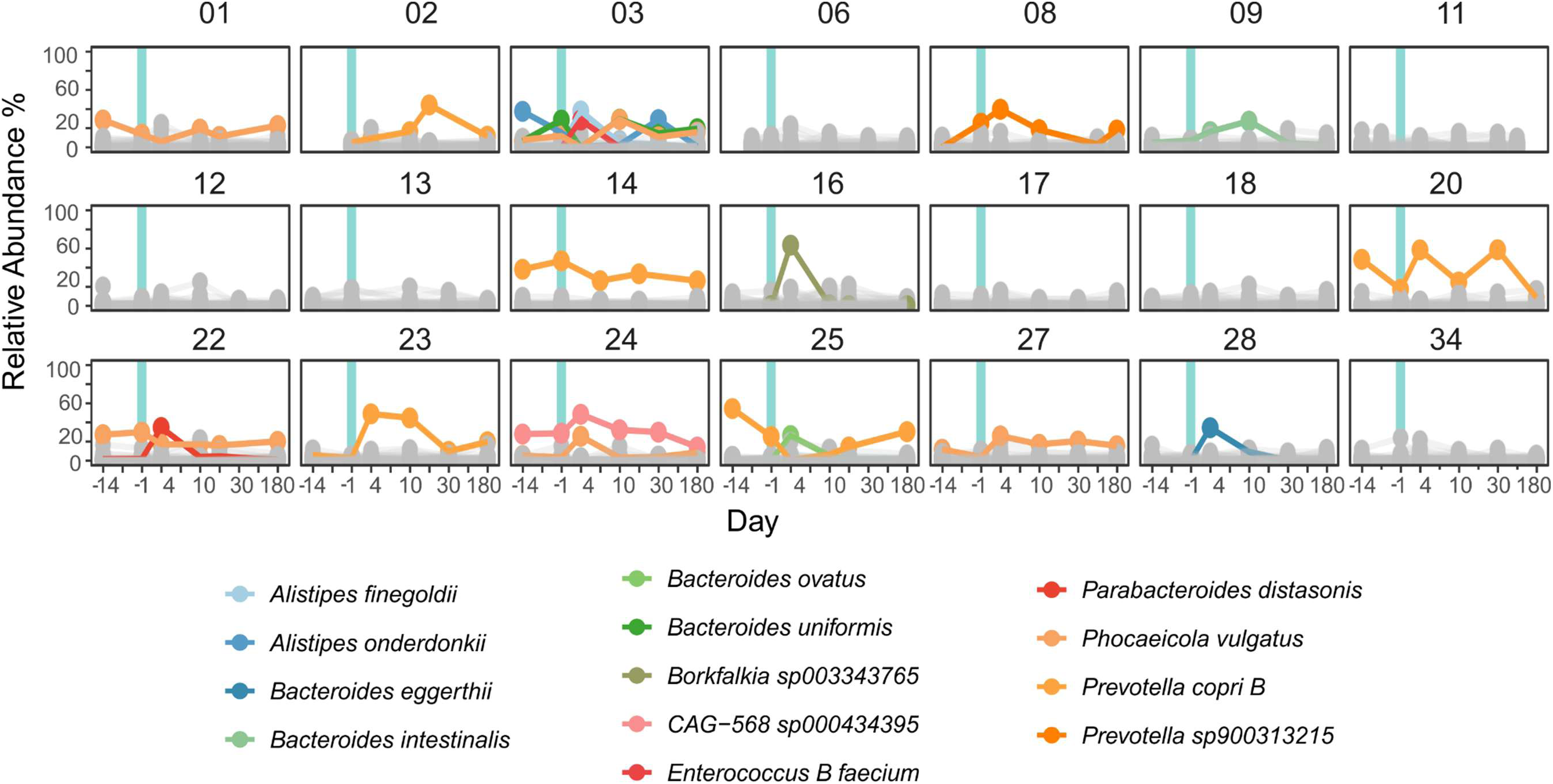
Relative abundances of bacterial gut species over the course of the clinical trial (as in Figure 4A for vOTUs).

## STAR Methods

### Key resources table

| REAGENT or RESOURCE | SOURCE | IDENTIFIER |
| --- | --- | --- |
| *Biological samples* |  |  |
| Fecal samples of healthy volunteers | d'Humières et al., (2024) | https://doi.org/10.1186/s40168-023-01746-0 and ClinicalTrials.gov ID: NCT02659033 |
| *Deposited data* |  |  |
| Shot gun sequencing data (bacterial fraction) | European Nucleotide Archive | PRJEB58157 |
| Shot gun sequencing data (viral fraction) | European Nucleotide Archive | PRJEB58815 |
| Rscripts and data tables | - | https://github.com/EpfeiferNutri/PrediRes.git<br>https://doi.org/10.1101/2025.02.07.636470 |
| *Chemicals and commercial assays* |  |  |
| PEG 8000 | Sigma-Aldrich | Saint-Louis, MO, USA |
| Sodium chloride (NaCl) | Sigma-Aldrich | Saint-Louis, MO, USA |
| PBS (Phosphate Buffered Saline) | Sigma-Aldrich | Saint-Louis, MO, USA |
| DNase | Sigma-Aldrich | Saint-Louis, MO, USA |
| Sephadex column | Sigma-Aldrich | Saint-Louis, MO, USA |
| SM buffer (100 mM NaCl, 8 mM MgSO<sub>4</sub>·7H<sub>2</sub>O, 50 mM Tris-Cl) | Lab-prepared | - |
| Phage DNA Extraction Kit | Norgen Biotek Corp | Thorold, ON, Canada |
| Qubit dsDNA HS Assay Kit | ThermoFisher Scientific | Waltham, MA, USA |
| Nextera XT Kit | Illumina | - |
| Illumina HiSeq 2500 PE_250 | Illumina | - |
| *Software, databases and algorithms* |  |  |
| Atropos (v1.1.18) | - | https://github.com/jdidion/atropos |
| Bowtie2 (v2.5.1) | - | http://bowtie-bio.sourceforge.net/bowtie2 |
| SPAdes (v3.15.5) | - | http://cab.spbu.ru/software/spades |
| BWA-MEM2 (v2.2.1) | - | https://github.com/bwa-mem2/bwa-mem2 |
| msamtools (v1.1.3) | Arumugam Lab | https://github.com/arumugamlab/msamtools |
| samtools (v1.4) | - | http://www.htslib.org/ |
| seqtk (v1.3) | - | https://github.com/lh3/seqtk |
| MUMmer (v4.0.0) | - | https://github.com/mummer4/mummer |
| Unified human gut virome database (UHGV) | - | https://github.com/snayfach/UHGV |
| Unified Human Gastrointestinal Genome catalogue (UHGG) (v2.0.2) |  | https://www.ebi.ac.uk/metagenomics/genome-catalogues/human-gut-v2-0-2 |
| geNomad (v1.7.4) | - | https://github.com/apcamargo/genomad |
| VIBRANT (v1.2.1) | - | https://github.com/AnantharamanLab/VIBRANT |
| viralVerify (v1.1.3) | - | https://github.com/ablab/viralVerify |
| CheckV (v1.0.1) | - | https://bitbucket.org/berkeleylab/checkv |
| Catalogue of 1416 phage-plasmids | Pfeifer and Rocha., 2024 | https://doi.org/10.1038/s41467-024-45757-3 |
| vConTACT2 (v0.11.3) | - | https://bitbucket.org/MAVERICLab/vcontact2 |
| blastn (v2.15.0) | - | https://hub.docker.com/r/ncbi/blast/tags |
| iPHoP (v1.3.3) | - | https://github.com/snayfach/iPHoP |
| BACPHLIP (v0.9.6) | - | https://github.com/adamhockenberry/bacphlip |
| AMRFinderPlus (v3.12.8) | - | https://github.com/ncbi/amr |
| METEOR (v1.0) | MetaGenoPolis | https://github.com/metagenopolis/meteor |
| Vegan R package (v2.6.4) | R Project | https://vegandevs.github.io/vegan/ |
| gggenomes (v1.0.1) | - | https://thackl.github.io/gggenomes/ |
| Gephi (v0.10.1) | - | https://gephi.org/ |

## Details to Methods

### Collection and Background of Fecal Samples

This study is based on a previous work^28^ of the CEREMI clinical trial. In particular, 22 healthy volunteers, both male and female, aged 18 to 65 years, with no exposure to antibiotics in the preceding 3 months or hospitalization in the last 6 months, participated in the study. In the clinical trial, donors received intravenous treatment with either 1 g ceftriaxone once daily (n=11) or 1 g cefotaxime three times a day (n=11) for three consecutive days. Fecal samples were collected starting from 15 days before to 180 days after treatment for most donors (detailed sampling scheme shown in Figure S1). For the majority (n=18 donors), six time points are available, for 3 donors five samples and for one donor with four samples, no time point before treatment was available. Only for donor 29, no samples prior to the treatment were available. For this reason, we used the read sets from donor 29 only for assemblies and their characterization, but excluded these sample from all ecological and abundance analysis.

### Preparation of virome DNA and sequencing

Viral particles were prepared as described in previous work^28,48^ from samples collected on days ranging from −14, to 180 days after treatment with the two antibiotics (Figure S1). Phages were precipitated following a polyethylene glycol (PEG) protocol as evaluated in a previous work^48^. Briefly, 1 gram of fecal sample was homogenized in 40 mL of phosphate-buffered saline (PBS) (Sigma-Aldrich, Saint-Louis, MO, USA), agitated with a mechanical laboratory agitator for 1 hour at 4 °C, centrifuged at 17,000 g for 5 min, and filtered at 2 µm and 0.45 µm. Phages were precipitated using PEG by adding 1 M solid NaCl and 10% (v/v) PEG 8000 (Sigma-Aldrich, Saint-Louis, MO, USA), and incubated overnight at 4 °C. Particles were pelleted (at 5,250 g for 1 hour at 4 °C), and re-suspended in 500 µL of SM buffer (NaCl 100mM, MgSO_4_.7xH2O 8mM, Tris-Cl 50mM). Bacterial DNA were digested by DNase (10 U/mL) (Sigma-Aldrich, Saint-Louis, MO, USA) for 30 min at 37 °C and then heated to 65 °C for 10 min to stop the reaction. DNA extraction was done using the “Phage DNA extraction” kit (Norgen Biotek Corp, Thorold, ON, Canada), purified on a Sephadex column (Sigma-Aldrich, Saint-Louis, MO, USA), quantified using the Qubit dsDNA HS Assay kit (ThermoFisher Scientific, Waltham, MA, USA), and sequenced using the Illumina HiSeq2500 PE_250 method with the Nextera XT kit and an input of 1 ng DNA.

### Read Processing, Assembly, and Curation of Multimeric Sequences

Illumina adapters and low-quality reads were removed using Atropos (v1.1.18)^67^ (trim -m 100 –q 20,20 – trim-n), and subsequently the reads were mapped against the human genome (GRCh38) with Bowtie2^68^ v.2.5.1 (--very-sensitive), retaining only high-quality non-human (HQ-nH) reads for downstream analysis. On average, 8.85 +/-3.04 Mio. reads were available (see Figure S4, Table S2).

Contigs were assembled by following three approaches:

1. HQ-nH reads were pooled per donor and assembled using SPAdes^69^ v.3.15.5 in metaviral mode (--- metaviral)^35^. Assemblies larger 3kb, were validated by mapping reads back to them using bwa-mem2^70^ v. 2.2.1 and keeping only alignments of good quality using msamtools v.1.1.3 (https://github.com/arumugamlab/msamtools) (filter -S -b -l 80 -p 95 -z 80).
2. Reads not matching were saved (with samtools^71^ 1.4 (samtools fastq -1 read.1.fq.gz -2 read.2.fq.gz -s/dev/null -o /dev/null -c 6 -N -) and mapped per sample to the unified human gut virome database (https://github.com/snayfach/UHGV) (UHGV) (https://portal.nersc.gov/UHGV/, last accessed on 19.01.2024). Viral operational taxonomic units (vOTUs) that were covered by at least 50% of sequence length with an average depth of 1 were retrieved.
3. In a few samples, more than 90% of reads did not align with any sequences obtained by the first two approaches and or to any further datasets (see Assessing Quality of Read Signal by Mapping). Of these low-covered samples, we randomly extracted reads (10% and 1%) using seqtk v.1.3 (https://github.com/lh3/seqtk). For 10%: sample -s100 read1.fq 0.1 > sub01.fq, and repeated the assembly with SPAdes^69^ v.3.15.5 (in default mode). The assemblies, referred as sub sampling approach, resulted in a higher final coverage (fractions shown in Figure S4).

Contigs from the three approaches were pooled, and only sequences >3kb were used to predict viral contigs (N=8966).

In addition, we screened these sequences for concatemeric forms using MUMmer^72^ 4.0.0 (exact-tandems contigs.fna 1000), and by reporting only sequences with tandems >1000 bp. Positive candidates were manually inspected and resolved to monomeric forms. In total, we found two of the total 8966 sequences as concatemeric.

### Defining viral contigs

Contigs were classed as viral if geNomad^34^ v.1.7.4 (end-to-end), VIBRANT^36^ v. 1.2.1 (VIBRANT_run.py), or viralVerify v.1.1^35^ (default) detected them as positive hits. Proviral sequences reported by geNomad and VIBRANT were added in a separate category (“proviral”) and not classed as viral. Intersection of the viral classification tools are shown in Figure S2.

### Clustering viral contigs into viral Operational Taxonomical Units

To define viral Operational Taxonomical Units (vOTUs) and curate the sequences from redundancy, we pooled all viral, assembled contigs with the ones detected in UHGV. We clustered sequences into vOTUs using a customized workflow. First, we computed the average nucleotide identity (ANI) between all sequences using blastn v.2.15.0^73^ and applied the anicalc.py script by CheckV^37^ 1.0.1 to compute the average nucleotide identity (ANI) between all contigs. Then, we constructed a network, where nodes are viral contigs and edges represent ANI values matching the viral species definitions (at least 95% ANI and 85% alignment fraction (AF) in reference to the shorter contig). Largest viral contigs were chosen as the representative sequences (vOTUs). Connected nodes were extracted as clusters. Singletons were directly considered as vOTUs. We observed that in some clusters some nodes were not directly connected to each other (belong to the same species), but by inter-connecting nodes which may bridge two related species level. To control for this, we performed an iterative curation of the clustered network: We first ranked contigs by size (largest first, since it is the representative) in ranges of 3 kb.

Then, all sequences related to the one with the highest rank were removed (from all clusters) from the network since they represent redundant sequences and belong to the same species. The elimination results in a new, less complex and connected, network. We then defined again connected nodes as clusters, repeat the elimination and curate step-wise for inter-connected sequences (from the largest to the smallest contigs) until the network consist of only of singletons with defined vOTUs. Overall, this approach resulted in 6467 non-redundant vOTUs (see Table S1).

### Taxonomical and functional annotation of viral contigs

We applied CheckV^37^ v.1.01 (with database version 1.5) to assess the quality and completeness of viral contigs. Taxonomy was predicted with geNomad^34^ and vConTACT v2^46^. geNomad gives directly a viral taxonomy in the output table. For vConTACT v2, we clustered 27 022 phage sequences (retrieved from GenBank (accessed in February 2024)) and 168536 viral sequences from UHGV with all 6467 vOTUs. If taxonomical assignments were inconsistent, we reported predictions from the two tools. For the vOTUs retrieved from UHGV, we kept the assigned taxonomy. In total, we assigned to 5378 of the 6467 vOTUs a host (class level) (Table S1). We used vConTACT v2^46^ to cluster all 6467 vOTUs into 409 viral clusters (VCs) with at least 3 members (Table S1). vConTACT v2 clusters species broadly at the genus level^46^. Outliers, overlaps and vOTUs assigned to ‘Clustered/Singleton’ were treated as singletons. iPHoP^74^ v.1.3.3 (database version September 2021) was applied to predict the bacterial hosts, assigning 4206 vOTUs. Notably, iPHoP is primarily trained for bacterial and archaeal viruses and exhibits a high false-positive rate for eukaryotic viruses^74^. It predicted a bacterial host for 27 out of the 31 eukaryotic vOTUs, consequently, we have low confidence in these predictions. BACPHLIP^40^ v.0.9.6 was applied on contigs of complete and high quality to predict viral life cycles. Antibiotic resistance genes were annotated by AMRfinderPlus^33^ v.3.12.8 on genes called by geNomad^34^.

### Clustering of vOTUs with phage-plasmids

We used our database of 1416 P-Ps to search for homologous sequences among the vOTUs. To achieve this, we grouped them by two approaches: 1) vConTACT v2^46^ (default) and 2) weighted gene repertoire relatedness (wGRR). wGRR was computed as described in previous works^16,32^. vOTUs with at least 10 genes and a relatedness of wGRR > 0.3 to a P-P were considered as potential P-Ps. Networks were visualized using Gephi^75^ and genome relations were visualized using gggenomes (https://thackl.github.io/gggenomes/). A brief comparison of the two methods showed that they work in a nearly complementary manner. While vConTACT v2 detected distant relationships that do not meet the wGRR >0.3 threshold, it tends to miss smaller genomes or genomes with only a few edges (despite strong homology). These cases are instead captured by the gene repertoire approach.

### Assessing Quality of Read Signal by Mapping

We evaluated assembly and read quality by mapping HQ-nH reads to different gut-related datasets (per sample) using bwa-mem2 with msamtools (as described above). Reads aligning to viral contigs were referred to as the viral signal. Non-mapping reads were sequentially mapped to non-viral contigs (assembled but not classed as viral), proviral sequences, UHGV, and UHGG^38^. The proportions of reads mapping to each dataset per sample and on average are shown in Figure S3 and Table S2.

### Quantitative and statistical analysis of vOTUs

To assess viral species abundance, we removed highly contaminated samples or those with insufficient viral signal (<10% viral signal and <1 million viral reads), removing 2 of 127 samples. We used bwa-mem2 for mapping and retained only reads with good-quality alignments using msamtools (filter -S -b -l 80 -p 95 -z 80). vOTUs were considered as present in a sample if the sequence was at least half covered with an average read depth of 1. To assess the diversity of samples, we computed the abundances of vOTUs. For this, we normalized the counts of reads mapping on a contig to the contig size, and rarefied (down-sized) these counts according to the minimal library size. For within-donor analyses (alpha diversity, relative abundance), rarefaction matched the smallest read count per donor. For beta-diversity, the smallest read count of all considered samples was chosen for the rarefaction. Alpha diversity was indexed by the Shannon and beta diversity by Bray-Curtis distance (with the vegan v2.6.4 package in R (https://vegandevs.github.io/vegan/)). Relative abundance was defined as the fraction of total abundance of all present contigs in a sample.

To assess phage population dynamics, we calculated the mean coefficient of variation (CV) for each vOTU within each donor. The CV, a measure of relative variability, was determined by dividing the standard deviation by the mean of relative abundance values. vOTUs absent from all samples of a donor were excluded from the analysis. For vOTUs present in at least one sample per donor, zero values were assigned to absent time points. The mean CV was calculated only for vOTUs with at least three values.

### Abundance profiling of metagenomic species

The abundances of bacterial species were retrieved from previous work^28^. Briefly, our catalogue of gut species is based on 2,741 unique metagenomic species. Microbial gene count table was created using METEOR (https://github.com/metagenopolis/meteor). Relative abundances of the species were calculated as the mean of their 100 signature genes, which show the highest correlation with each other (per species). If fewer than 10% of these genes were detected, the abundance was set to 0. Relative abundance was computed by normalizing to the sum of all species to estimate the proportion. Bacterial richness in each sample was defined as the number of unique species. An overview of the microbial species is listed in Table S5 and their relative abundances in Table S6. To assess fluctuations of species per donor, we computed the CV as described for the vOTUs.

### Goodness-of fit test for the number of dominant phages on distinct days

To test for significant differences in the number of highly abundant vOTUs on distinct days, we used a Poisson regression model^76^, by setting day before treatment (day -1) as the baseline, and considering the number of samples per day. We used the glm() function of the stats package in R to model the Poisson distribution with ‘family = poisson(link = “log”)’ and ‘offset = log (n_samples_per_day)’. We considered only days with more than three samples. P-values for all days were computed for different thresholds of dominant vOTUs starting from 0.1% to 40.0% (Figure S8BC).

