## supplementary file 1 for "Antibiotic perturbation of the human gut phageome preserves its individuality and promotes blooms of virulent phages"

Supplemental file 1.

Slide 2: Comparison of detection methods and lists of vOTUs distinct for each method.

Slide 3: Networks of vOTUs and P-Ps based on genome relatedness scores. Genome similarity was computed using protein-sharing networks. P-Ps and vOTUs (nodes) were grouped using the wGRR metric (edges). Only edges >0.3 are shown.

Slide 4: vContact v2 was used to cluster P-Ps and vOTUs (nodes). Edges are weights from the vContact v2 generated network metric.

Slide 5-32: Genome-to-genome synteny plots generated with gggenomes (https://github.com/thackl/gggenomes). Gene-to-gene assignments are the best-bidirectional-hits as used to compute the wGRR. RefSeq accession number are indicating P-P sequences from PMID: 38378896.

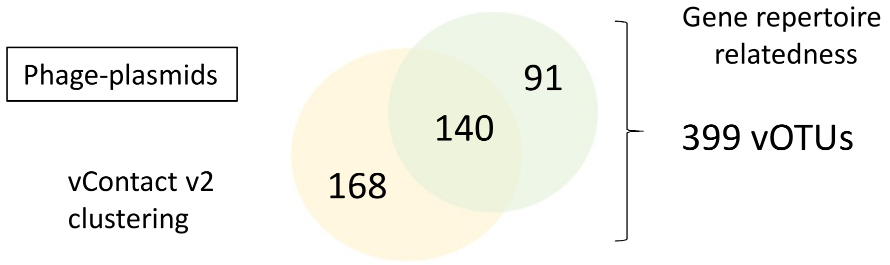


vContact v2 vs wGRR (>0.3) relatedness

**N = 168 distinct**

N = 61 related *Enterococcus faecium* p63-3, 18 kb
and *Clostridium beijerinckii* plasmid 18 kb
N = 41 related to a P-P of *Corynebacterium atypicum*;
were not detected by wGRR since far related (<0.3)
N = 32 related to vB_CpeS-CP51, similar Rep as *Carjivirus communis*

N = 8 related to CTC plasmid
N = 5 x *Selenomonas ruminantium lactilytica*, plasmid, 35 kb
N = 5 x Sarcina sp. (Clostridiaceae) plasmid p2 34 kb
N = 5 x related to *Carjivirus communis* (far related)

N = 3 x P-Ps of pBS32
N = 3 x *Cupriavidus oxalaticus, plasmid 38 kb*
N = 2 x *Klebsiella pneumoniae* plasmid, 40 kb
N = 2 x two P-Ps of IEBH
N = 1 x six PPs of pLP39

**N = 91 distinct**

N = 72 related to CTC plasmid,
N = 5 related to *Carjivirus communis* (fragments)
N = 3 related to SSU5 (super) group

N = 2 related to a P-P *of Corynebacterium atypicum*
N = 2 x N15 (1xgroup,1xcomm)
N = 2 x F116 (community)
N = 2 x *Enterococcus faecium* plasmid, 58 kb

N = 1 x *Escherichia coli* pEM03-18-08_2, 39 kb
N = 1 x *Lactococcus lactis* plas2, 35 kb
N = 1 x *Pseudomonas luteola* plasmid, 120 kb

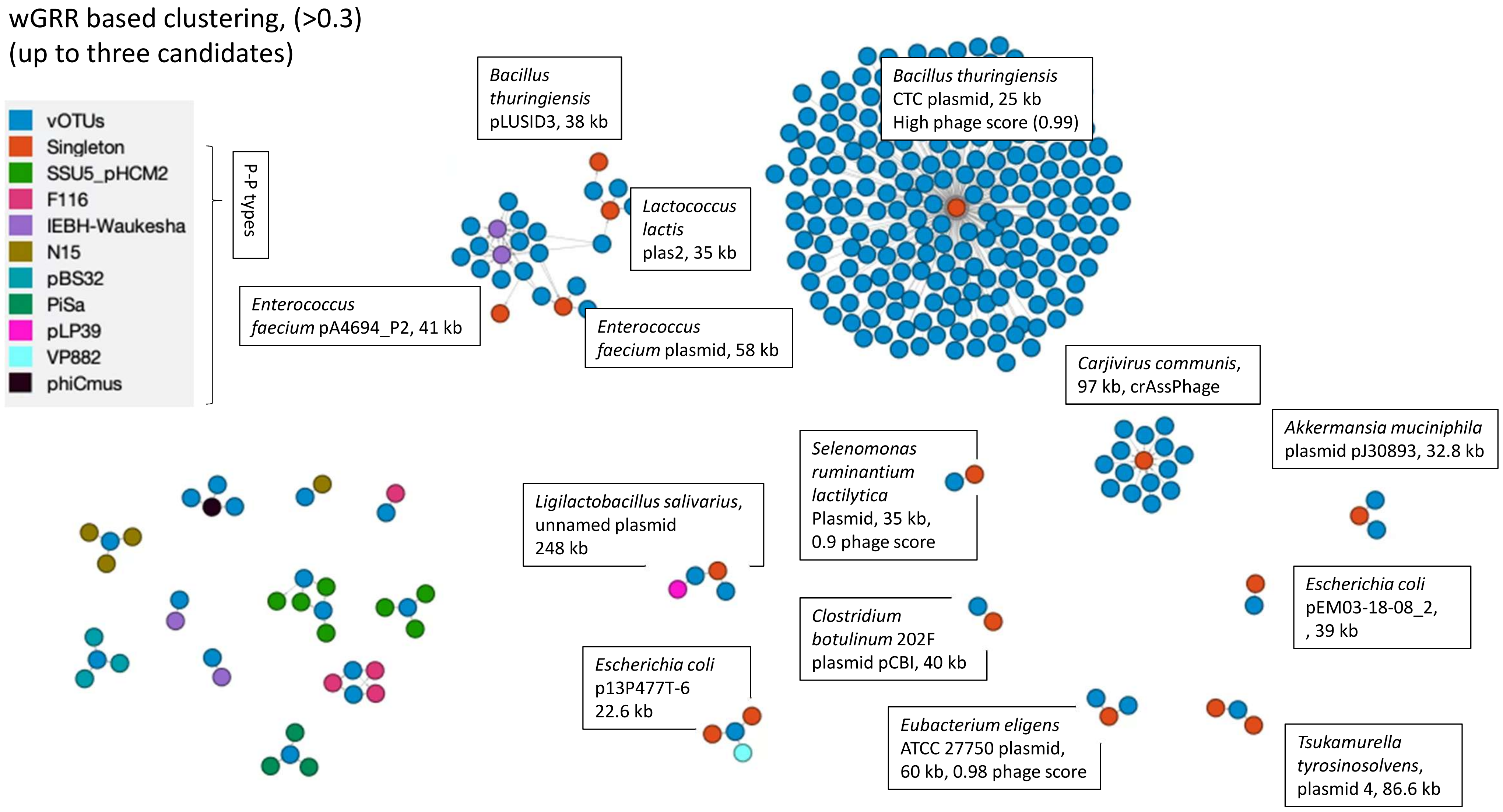

wGRR based clustering, (>0.3)
(up to three candidates)
vOTUs
Singleton
SSU5_pHCM2
F116
IEBH-Waukesha
N15
pBS32
PiSa
pLP39
VP882
phiCmus
P-P types
Bacillus thuringiensis pLUSID3, 38 kb
Bacillus thuringiensis CTC plasmid, 25 kb High phage score (0.99)
Lactococcus lactis plas2, 35 kb
Enterococcus faecium pA4694_P2, 41 kb
Enterococcus faecium plasmid, 58 kb
Carjivirus communis, 97 kb, crAssPhage
Akkermansia muciniphila plasmid pJ30893, 32.8 kb
Selenomonas ruminantium lactilytica Plasmid, 35 kb, 0.9 phage score
Ligilactobacillus salivarius, unnamed plasmid 248 kb
Escherichia coli pEM03-18-08_2, , 39 kb
Clostridium botulinum 202F plasmid pCBI, 40 kb
Escherichia coli p13P477T-6 22.6 kb
Eubacterium eligens ATCC 27750 plasmid, 60 kb, 0.98 phage score
Tsukamurella tyrosinosolvens, plasmid 4, 86.6 kb

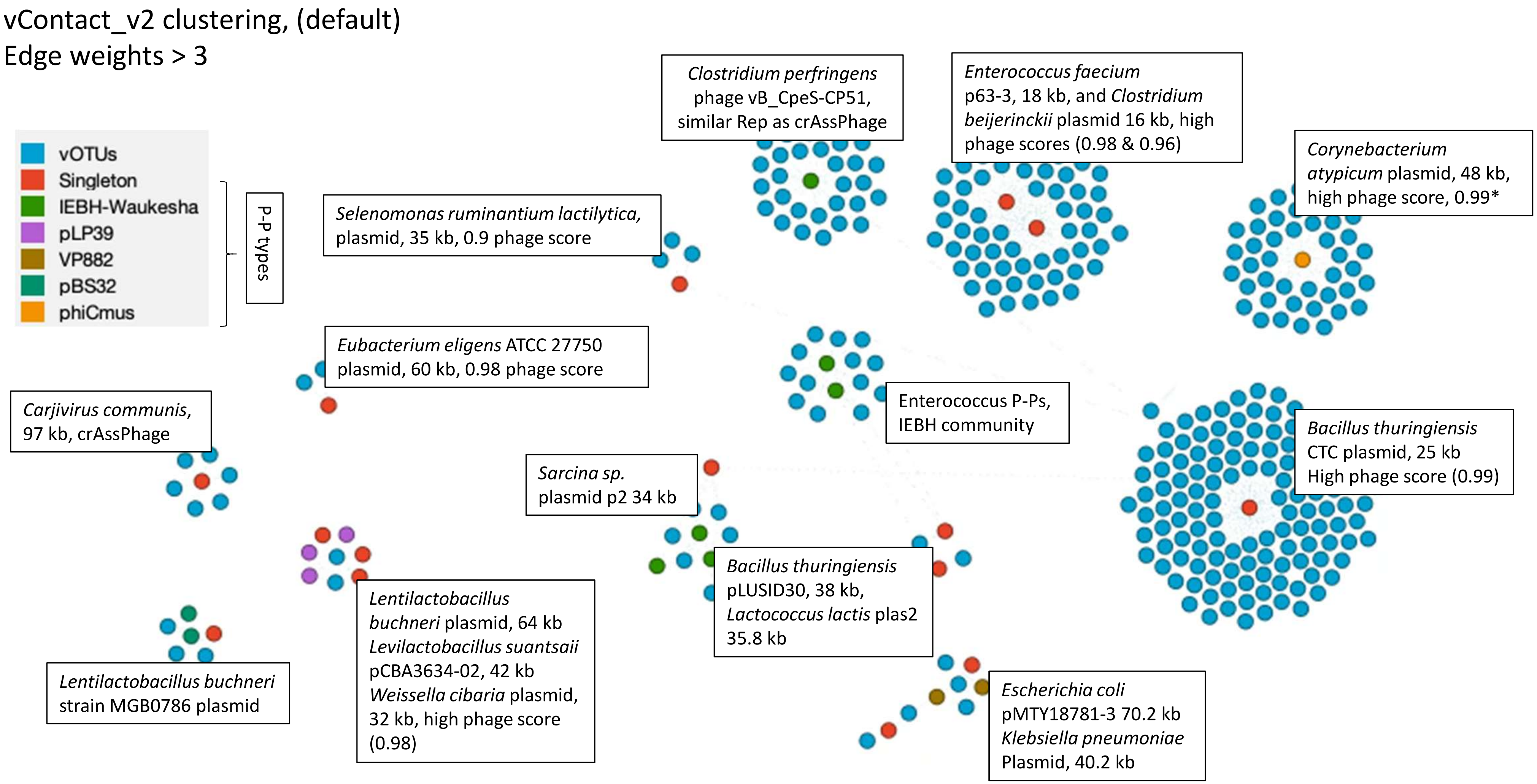
vContact_v2 clustering, (default)
Edge weights > 3
vOTUs
Singleton
IEBH-Waukesha
pLP39
VP882
pBS32
phiCmus
P-P types
Clostridium perfringens phage vB_CpeS-CP51, similar Rep as crAssPhage
Enterococcus faecium p63-3, 18 kb, and Clostridium beijerinckii plasmid 16 kb, high phage scores (0.98 & 0.96)
Corynebacterium atypicum plasmid, 48 kb, high phage score, 0.99*
Selenomonas ruminantium lactilytica, plasmid, 35 kb, 0.9 phage score
Eubacterium eligens ATCC 27750 plasmid, 60 kb, 0.98 phage score
Enterococcus P-Ps, IEBH community
Carjivirus communis, 97 kb, crAssPhage
Bacillus thuringiensis CTC plasmid, 25 kb High phage score (0.99)
Sarcina sp. plasmid p2 34 kb
Bacillus thuringiensis pLUSID30, 38 kb, Lactococcus lactis plas2 35.8 kb
Lentilactobacillus buchneri plasmid, 64 kb Levilactobacillus suantsaii pCBA3634-02, 42 kb Weissella cibaria plasmid, 32 kb, high phage score (0.98)
Lentilactobacillus buchneri strain MGB0786 plasmid
Escherichia coli pMTY18781-3 70.2 kb Klebsiella pneumoniae Plasmid, 40.2 kb

Detection method,
vContact v2 = 8
wGRR + vcontact = 118
wGRR = 72

Bacillus thuringiensis strain CTC plasmid

wgrr

0.41

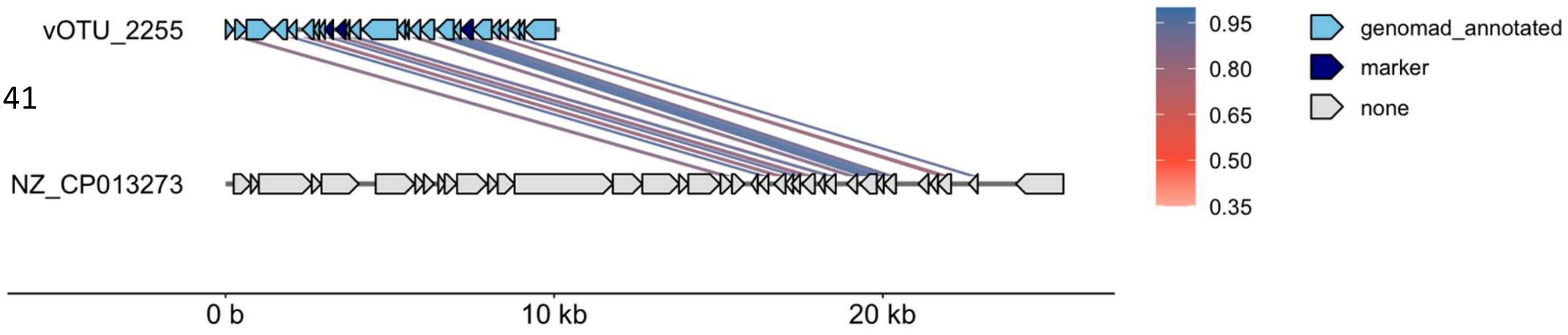

Detection method,
vContact v2 = 61

Enterococcus faecium isolate 2014-VREF-63 plasmid p63-3 sequence.

wgrr

0.11

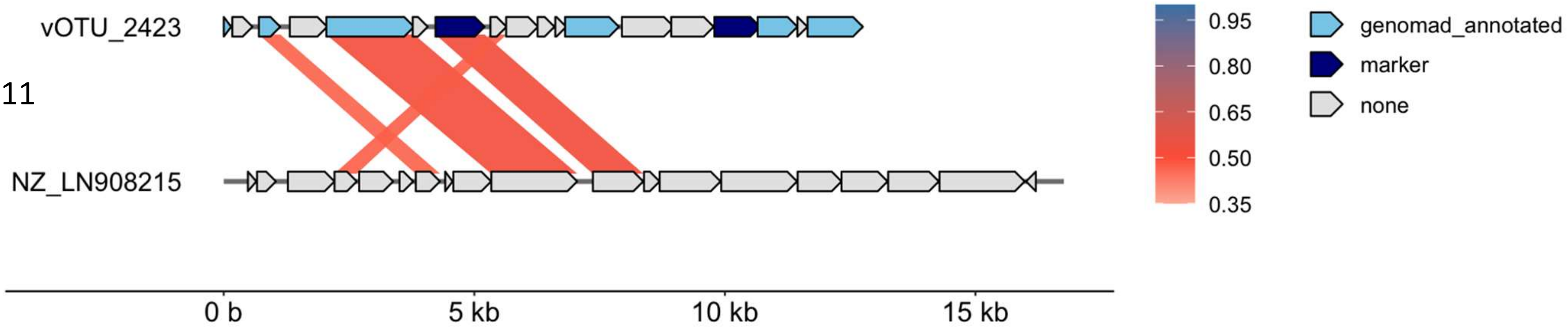

Detection method,
vContact v2 = 41
vContact v2+wGRR = 1
wGRR = 2

Corynebacterium atypicum strain R2070 plasmid phiCATYP2070I

wgrr

0.33

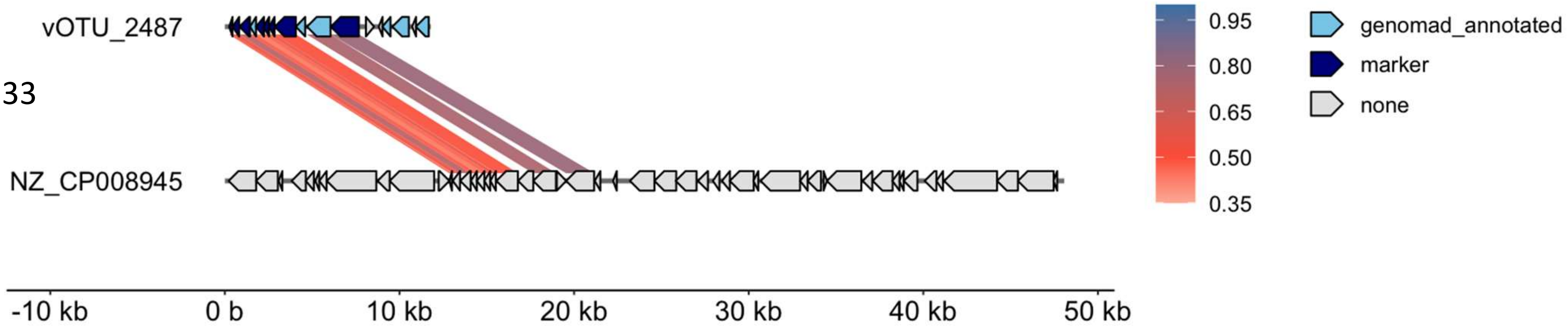

Detection method,
vContact v2 = 32

Clostridium phage vB_CpeS-CP51

wgrr

0.08

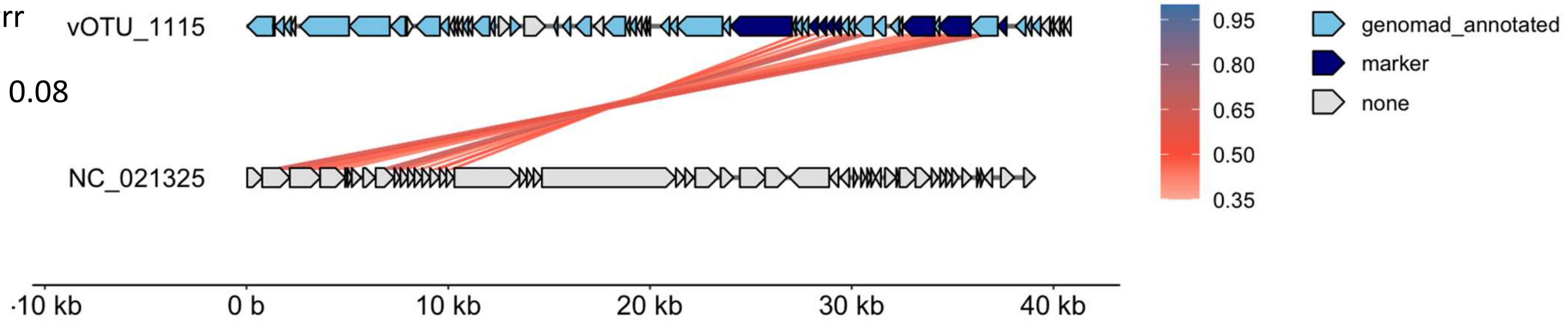

Detection method,
vContact v2+wGRR = 13

Enterococcus faecium isolate EFE11651 genome assembly, plasmid

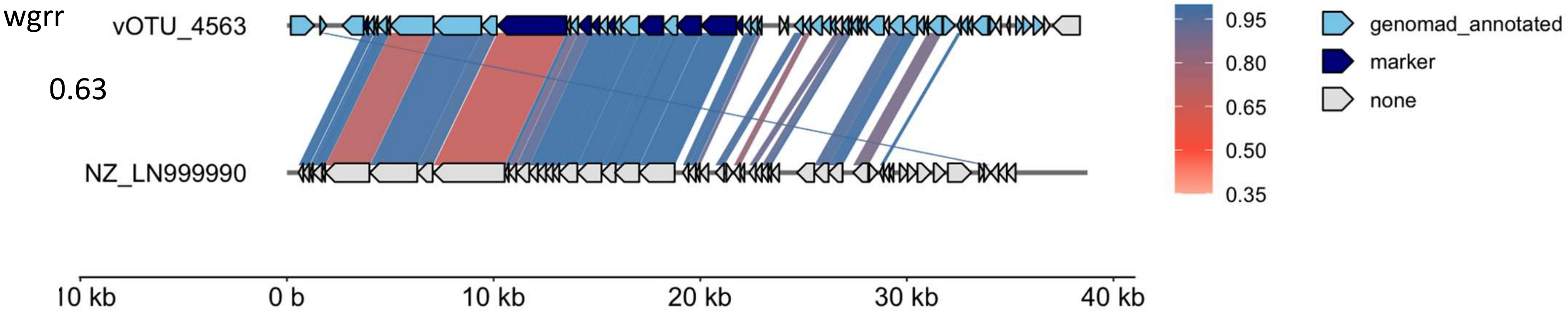

Detection method,
vContact v2 = 5
vContact v2+wGRR = 1
wGRR = 5

Carjivirus communis

wgrr

0.86

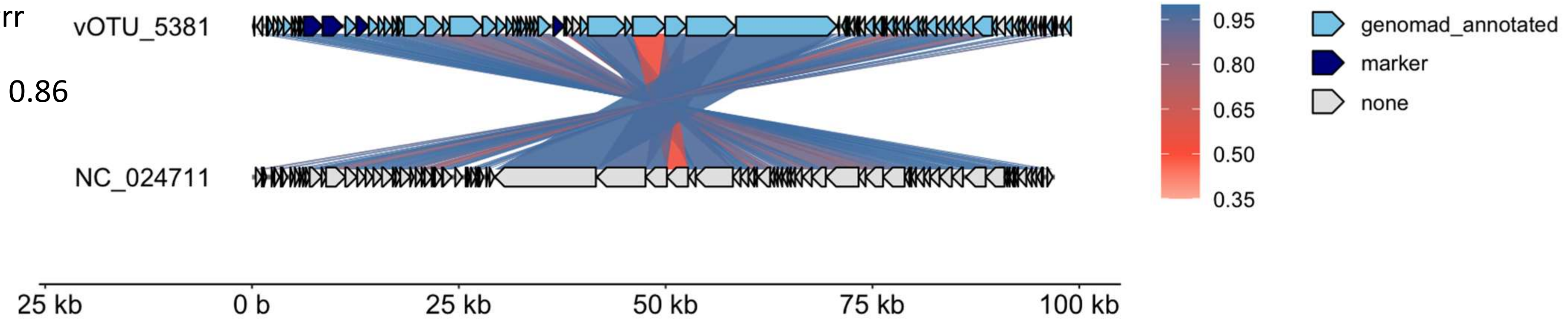

Detection method,
vContact v2 = 5

Selenomonas ruminantium subsp. lactilytica TAM6421 plasmid pSRC5 DNA

wgrr

0.14

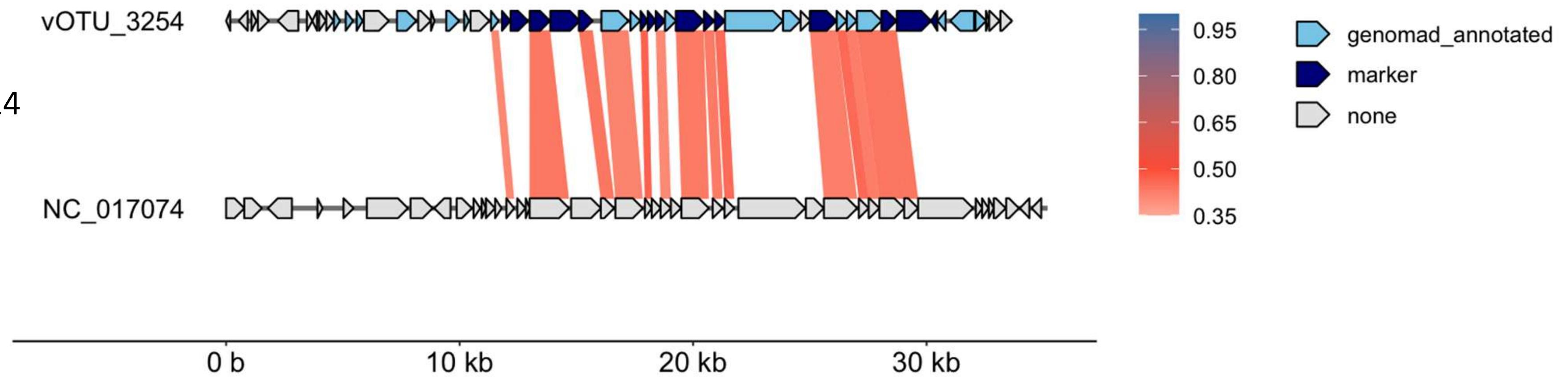

Detection method,
vContact v2 = 5

Paeniclostridium sordellii strain AM370 plasmid pRSJ16_1

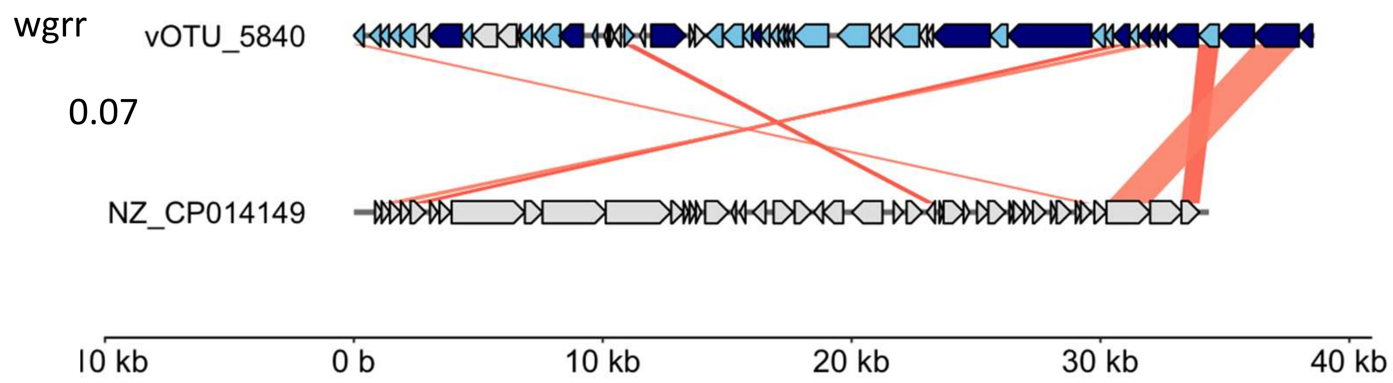

Detection method,
vContact v2 = 3

Arsenophonus nasoniae strain FIN plasmid pArsFIN14

wgrr

0.07

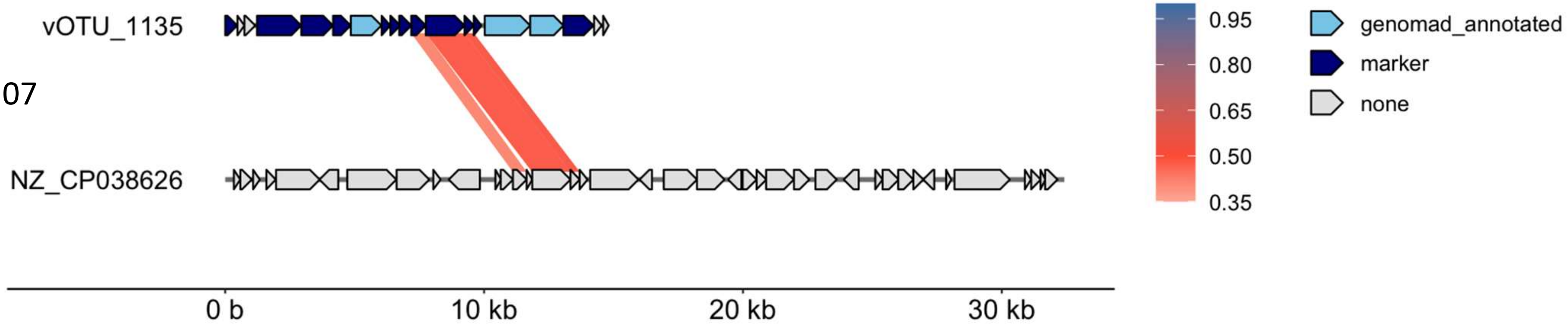

Detection method,
vContact v2 = 3

Bacillus licheniformis strain TAB7 plasmid pTAB7B

wgrr

0.15

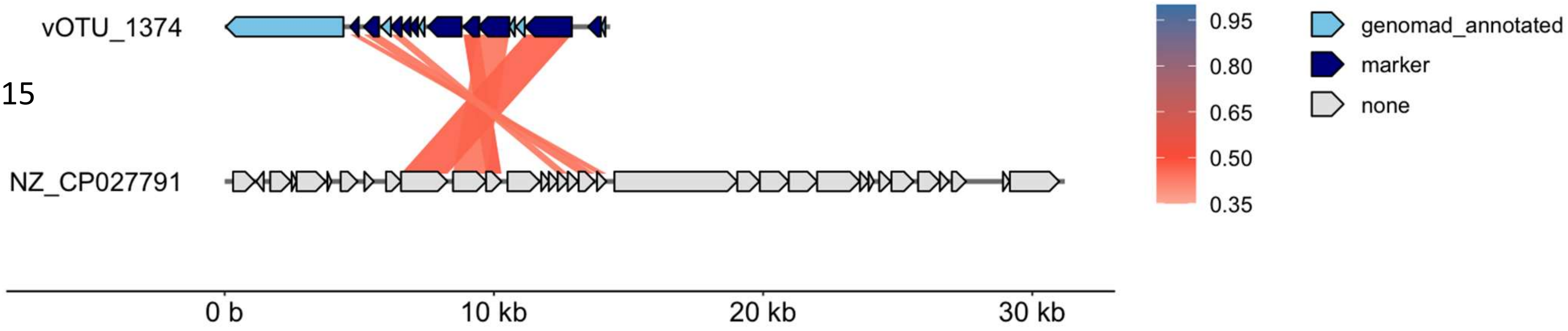

Detection method,
vContact v2+wGRR = 2

Akkermansia muciniphila plasmid pJ30893

wgrr

0.65

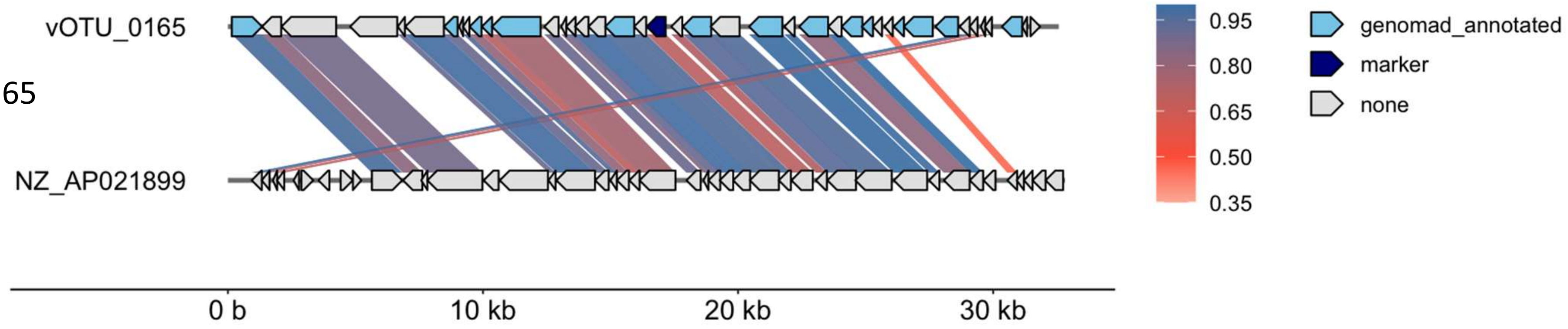

Detection method,
wGRR = 2

Enterococcus_faecium_strain_FA3_plasmid_unnamed2

wgrr

0.52

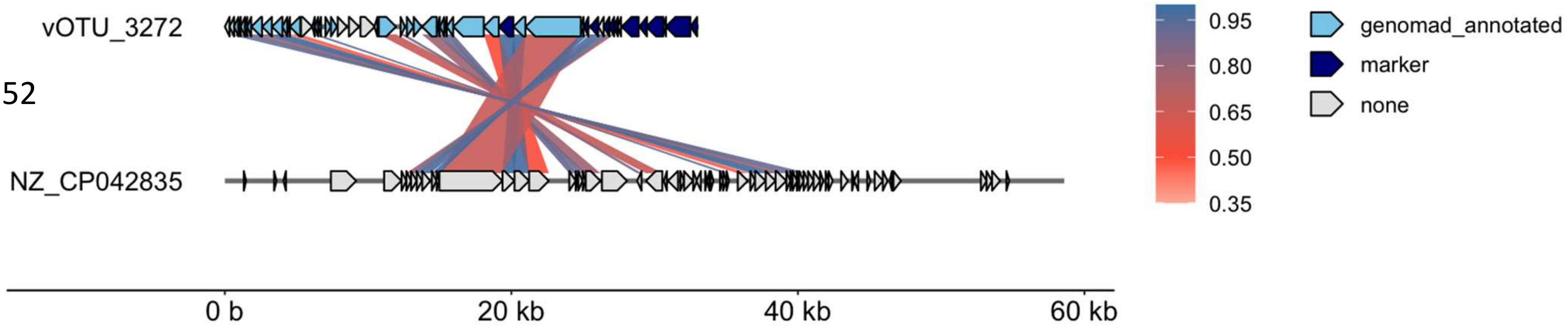

Detection method,
vContact v2+wGRR = 2

Eubacterium eligens ATCC 27750 plasmid unnamed

wgrr

0.34

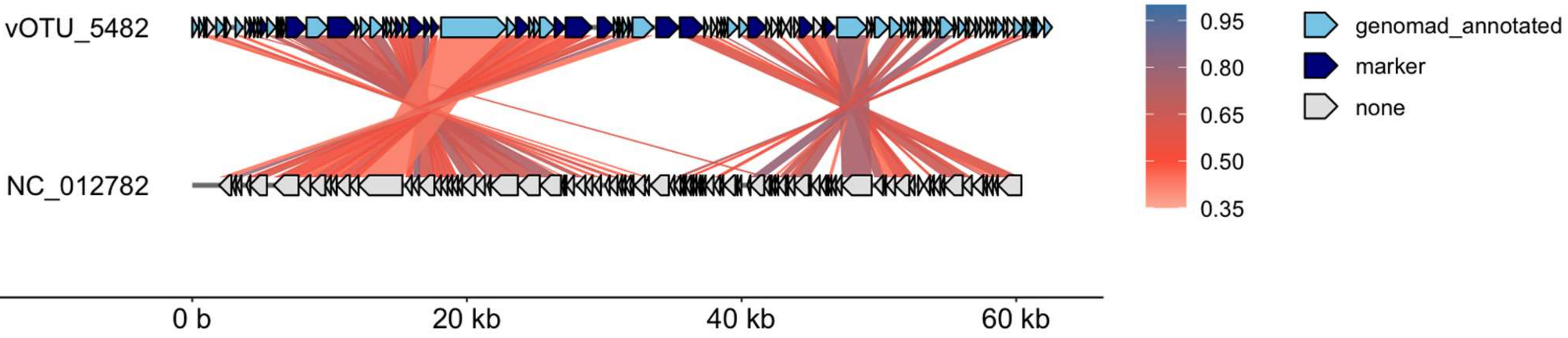

Detection method,
vContact v2 = 2

Clostridium phage PhiS63

wgrr

0.07

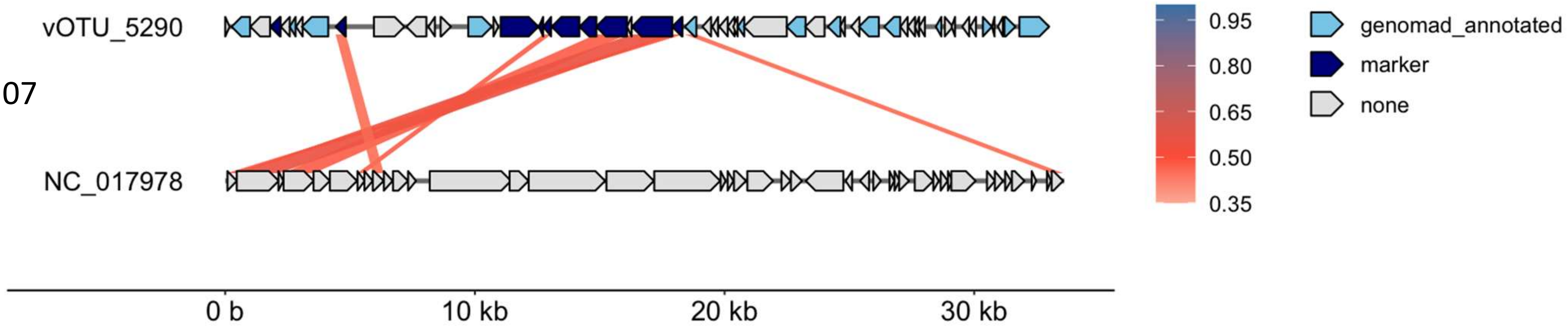

Detection method,
wGRR+vcontact = 2

Lactococcus lactis subsp. lactis strain L19 plasmid plas2, complete sequence

wgrr

0.35

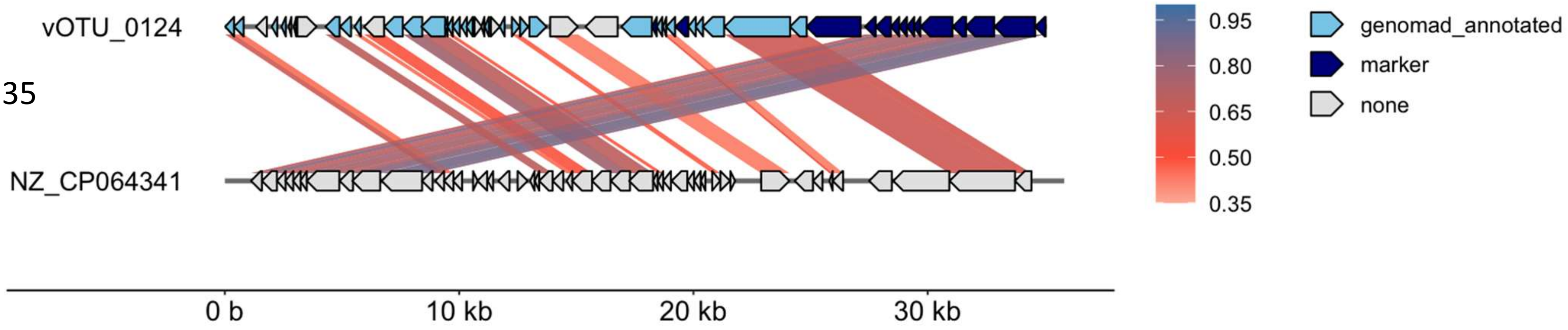

Detection method,
vContact v2 = 2

Klebsiella_pneumoniae_isolate_INF168-sc-2280023_plasmid_3,_complete

wgrr

0.18

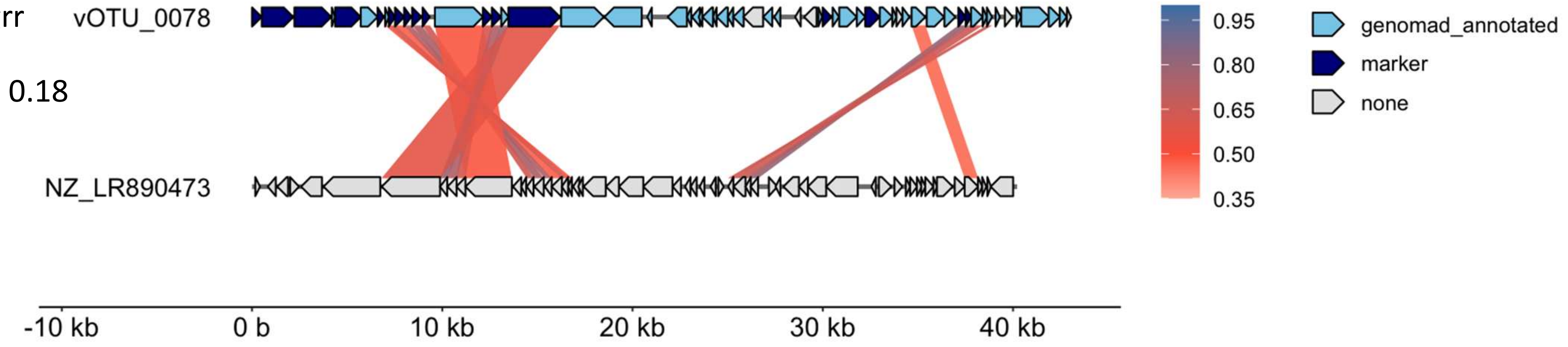

Detection method,
vContact v2+wGRR = 1
vContact v2 = 1

Lactobacillus reuteri I5007 plasmid pLRI02

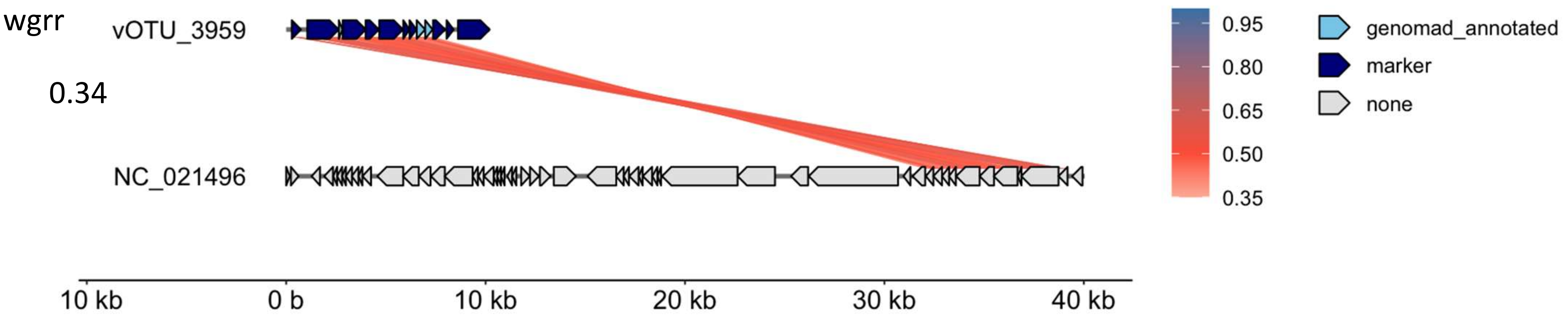

Detection method,
wGRR = 1

Citrobacter_sp._RHBSTW-00570_plasmid_pRHBSTW-00570_3,_complete

wgrr

0.91

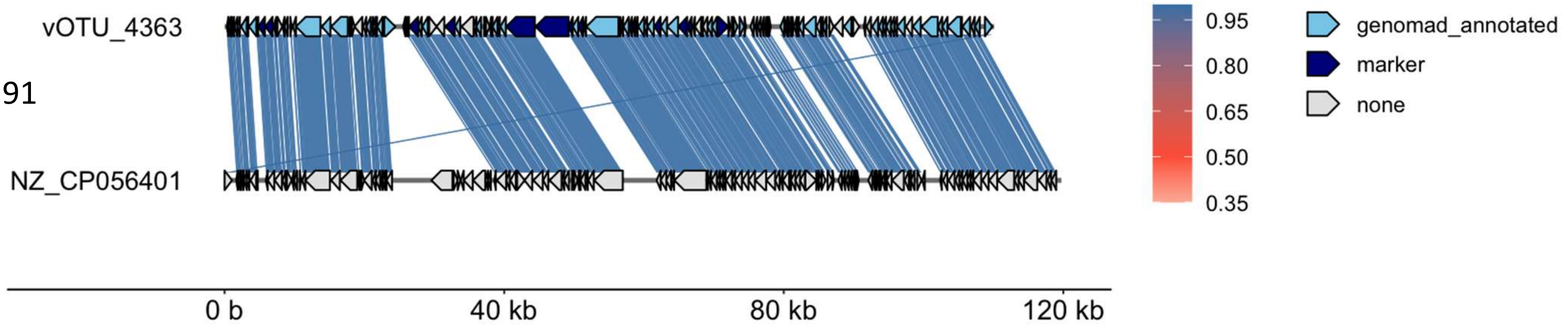

Detection method,
wGRR = 1

Klebsiella pneumoniae strain KPNIH36 plasmid pKPN-fff

wgrr

0.51

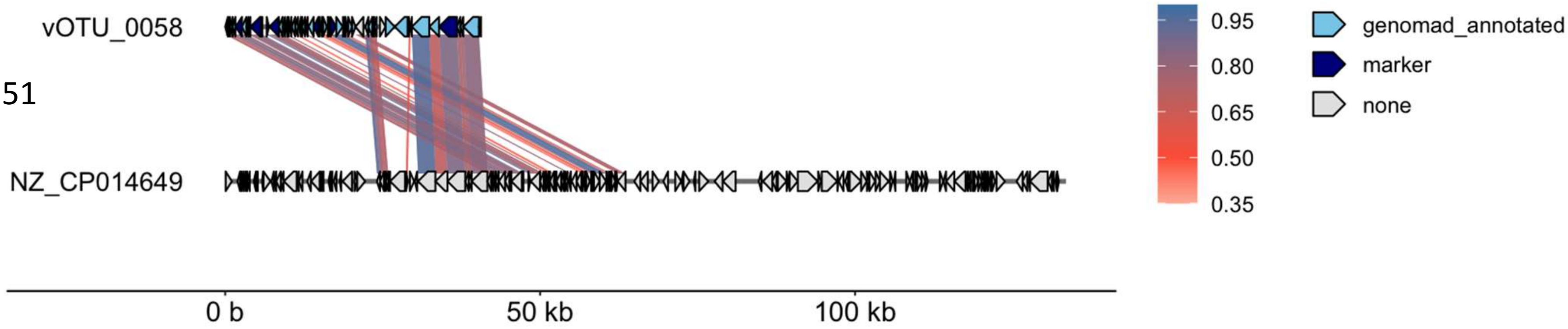

Detection method,
wGRR = 1

Escherichia_coli_strain_13TMH22_plasmid_p13TMH22-2,_complete

wgrr

0.66

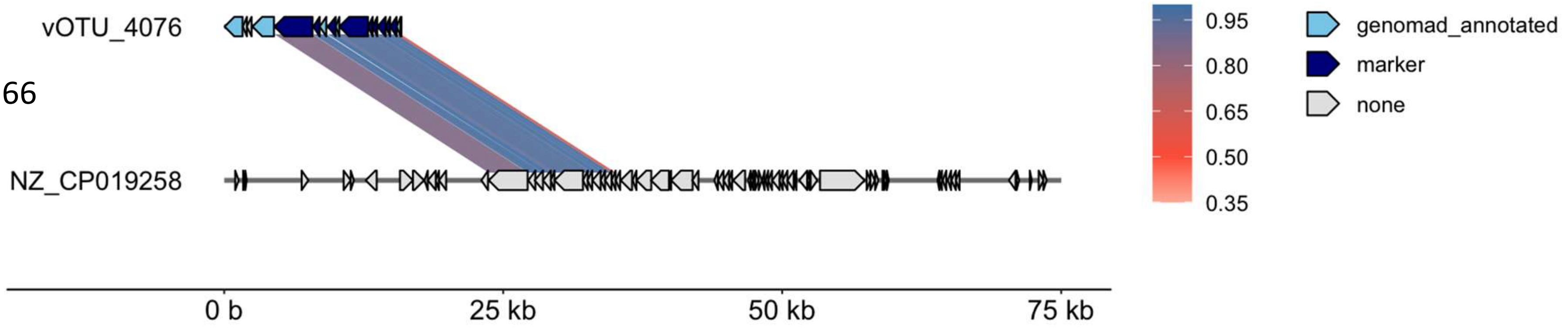

Detection method,
wGRR = 1

Enterobacter_hormaechei_strain_E70_plasmid_pE70

wgrr

0.91

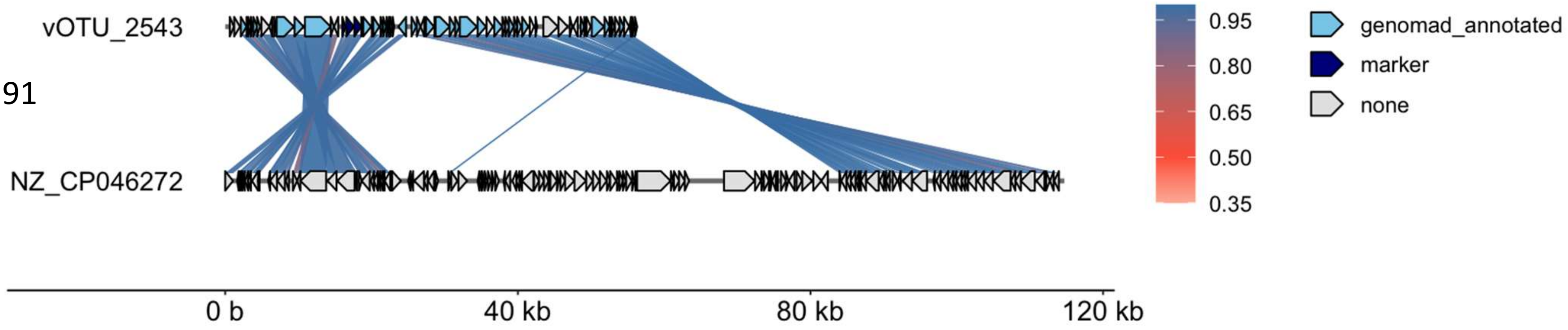

Detection method,
wGRR = 1

Klebsiella oxytoca strain KONIH2 plasmid pKOR-0e8e

wgrr

0.35

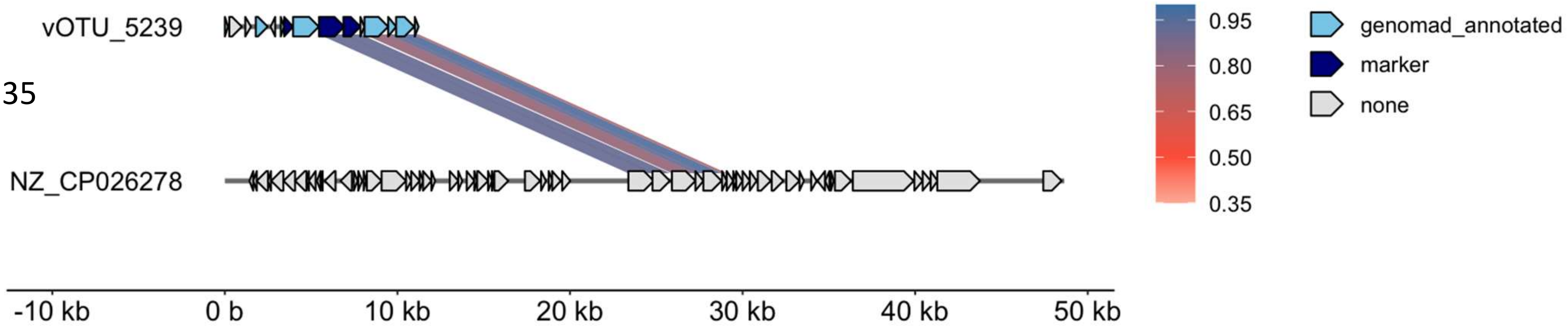

Detection method,
wGRR = 1

Klebsiella pneumoniae strain KPNIH36 plasmid pKPN-fff

wgrr

0.51

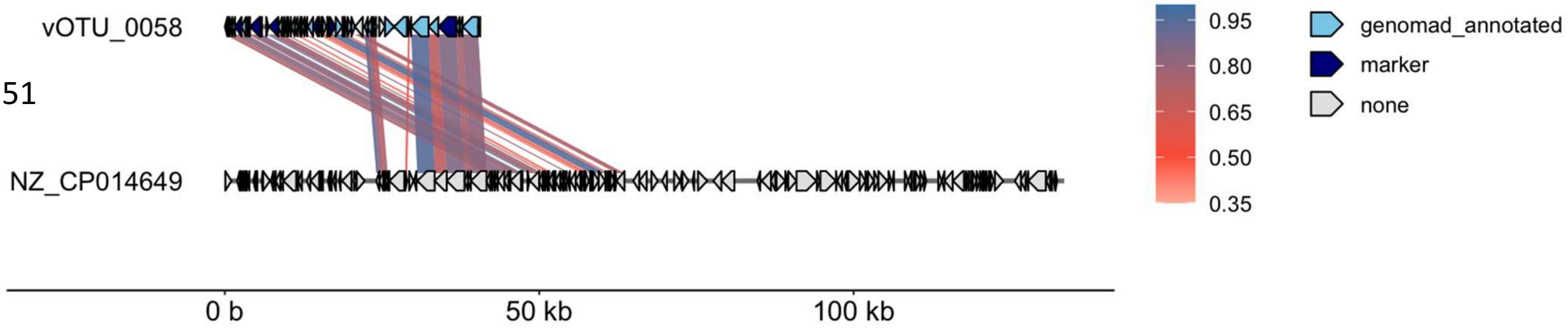

Detection method,
wGRR = 1

Enterobacter_hormaechei_strain_189_plasmid_pECL189-2,_complete

wgrr

0.79

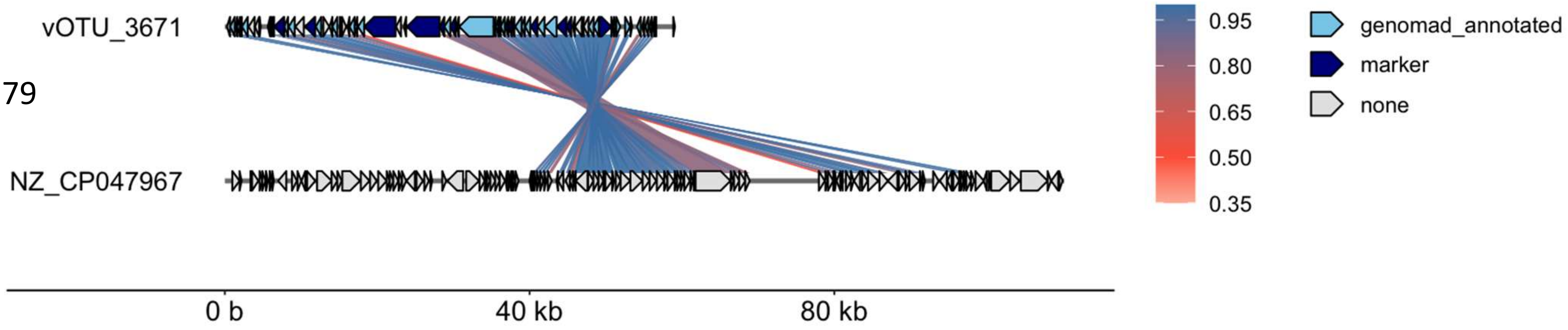

Detection method,
wGRR = 1

Escherichia_coli_strain_EM03-18-08_plasmid_pEM03-18-08_2,_complete

wgrr

0.32

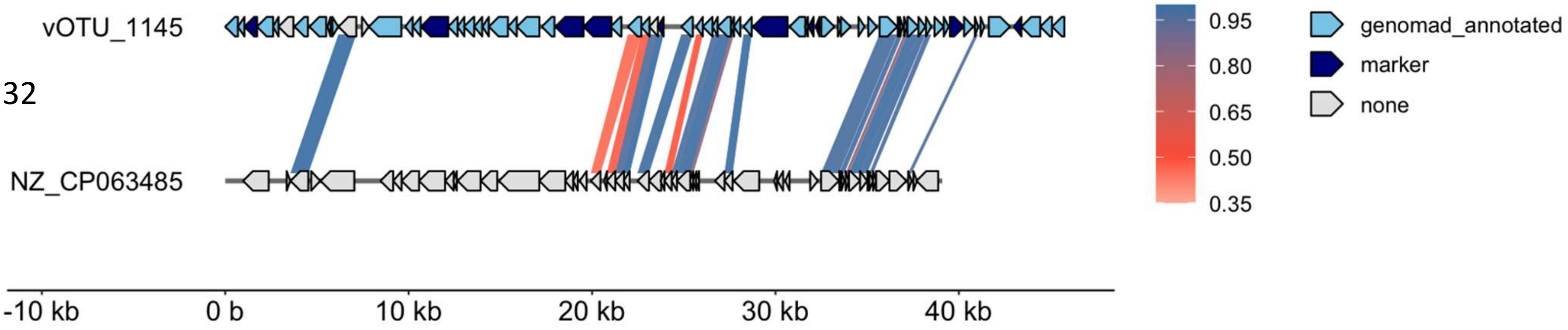

Detection method,
wGRR = 1

Escherichia_coli_O145_strain_RM9154-C1_plasmid_p1RM9154-C1

wgrr

0.36

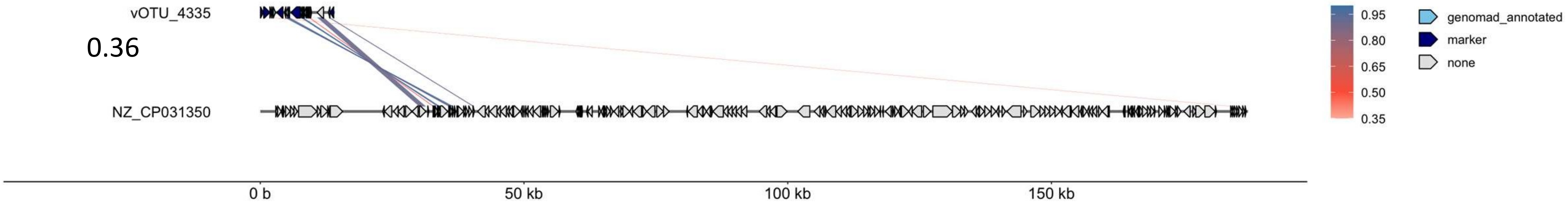

Detection method,
wGRR = 1

Lactococcus_lactis_subsp._lactis_strain_L19_plasmid_plas2,_complete

wgrr

0.65

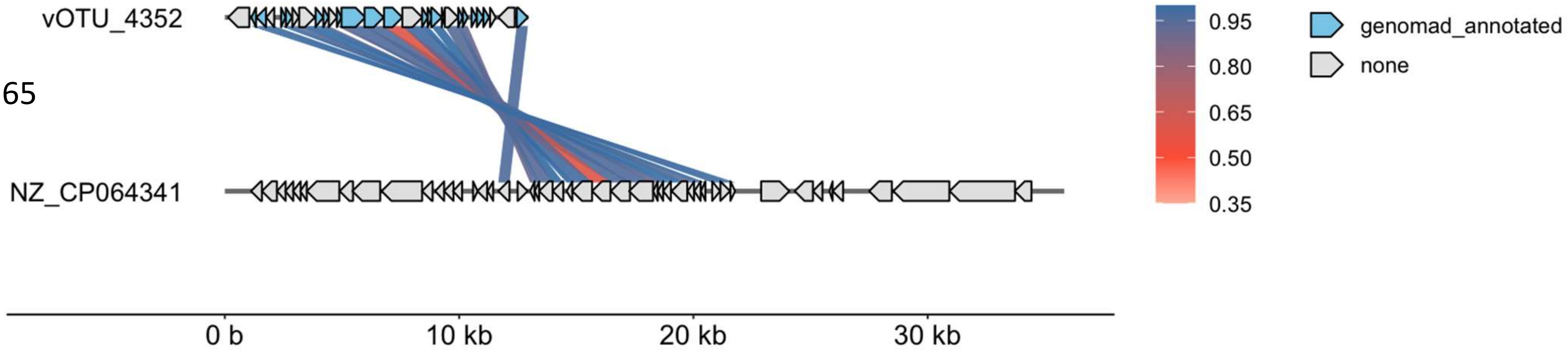

Detection method,
wGRR = 1

Pseudomonas_luteola_strain_FDAARGOS_637_plasmid_unnamed2

wgrr

0.45

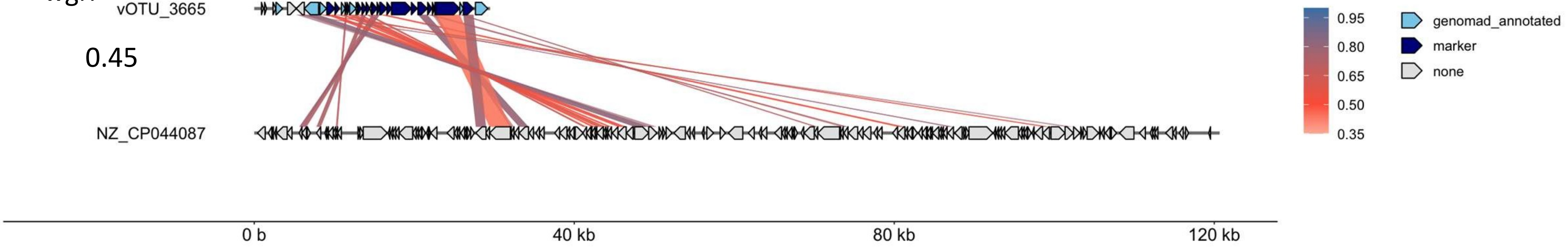
